# The role of medicinal Cu^2+^ salts in expanding the multi-pathway application of tigecycline against drug-resistant bacteria

**DOI:** 10.1101/2025.10.09.681327

**Authors:** Jinjing Xue, Qianyu Zhou, Yuanyuan Liu, Lin Ma, Houru Liu, Qianghua Lv, Kuan Gu, Jianfeng Wang, Yonglin Zhou

## Abstract

Tetracyclines are highly susceptible to photolysis, causing pollution in water, land and other ecosystems, as well as posing threats to human health. In this study, we found that only tigecycline and minocycline reduce the antibacterial activity in light, and this inferiority has severely hindered their scope of clinical use. By strategically regulating the photolysis pathway, we identified chalcanthite, which primarily contains Cu^2+^. By conducting antibiotic susceptibility test and electron microscopy analysis, Cu^2+^ salts (including copper sulfate, copper chloride, and copper gluconate) were confirmed to restore the antibacterial activity of tigecycline under light exposure against all tested bacterial strains. Spectroscopic characterization combined with quantum chemical calculations elucidated the molecular mechanism by which Cu^2+^ selectively modulates the photolytic degradation pathway of tigecycline and structural determinants (dimethylamino groups on the D ring). Multi-omics technologies revealed the mechanism underlying the Cu^2+^-mediated (non-antibacterial) inhibition of bacterial ferric citrate transporter. Animal infection models were developed using diverse drug-resistant pathogens to validate the *in vivo* therapeutic efficacy of copper gluconate-tigecycline combination therapy under photic conditions. These findings revealed effective adjuvants that facilitate the development of topical tigecycline formulations, and the establishment of a scientific foundation for synthesizing photostable tetracycline-based antibiotics through structural optimization.

## 1. Introduction

Tigecycline, a last-resort antibiotic, has experienced a continuous increase in global demand due to the emergence and rapid dissemination of multidrug-resistant pathogens [1–6]. Regarded as a cornerstone broad-spectrum antibiotic in clinical practice, Tigecycline is extensively prescribed for severe infections, including complicated intra-abdominal infections, skin and soft tissue infections, and community-acquired pneumonia [1–3, 5]. Tigecycline has a strong antibacterial effect on almost all gram-positive bacteria, gram-negative bacteria, and some anaerobic bacteria, especially pathogens resistant to traditional antibiotics [2–4]. High-level resistance of gram-negative bacteria to tigecycline and next-generation tetracyclines is mainly mediated by the tetracycline-inactivating enzyme Tet(Xs) [7–10]. Importantly, the antibiotic restriction policies in livestock farming has substantially mitigated the clinical threat posed by Tet(X4)-mediated resistance. Thus, tigecycline has long-lasting therapeutic value and significant developmental potential, supported by its preserved efficacy against all extensively drug-resistant (XDR) gram-negative bacterial strains except for *Pseudomonas aeruginosa* (*P. aeruginosa*).

As tetracyclines are extensively used, the toxicity of their parent compounds and degradation products in biological systems and ecosystems has been comprehensively investigated, and the mechanism underlying their photolytic propensity in aqueous environments has been well-elucidated [11–16]. Although the broad-spectrum antibacterial activity of tigecycline, a minocycline derivative, is greater than that of conventional tetracyclines, it shares inherent photosensitivity [15, 16]. This critical instability necessitates stringent light avoidance protocols during storage, transportation, and clinical administration, thereby restricting its therapeutic application for skin and soft tissue infections to injectable formulations. With recent advancements in medicinal chemistry, researchers have identified photostable products, represented by eravacycline and sarecycline, which mitigate degradation through structural optimization [17–19]. Resolving the disadvantage of easy photolysis of tigecycline can facilitate expanded clinical administration routes while achieving substantial resource optimization in pharmaceutical logistics.

Our preliminary investigations revealed divergent photolysis-activity relationships among tetracyclines: conventional analogs such as tetracycline, aureomycin, doxycycline and oxytetracycline retained measurable antibacterial efficacy post-photolysis, whereas tigecycline and minocycline exhibited light intensity-dependent attenuation of activity. To address this photochemical vulnerability, we implemented a high-throughput screening strategy from a traditional mineral pharmacopeia library. We reported that *Chalcanthite* (CuSO₄·5H₂O), a mineral-derived traditional Chinese medicine, restored the antibacterial efficacy of tigecycline under light-exposed conditions, and confirmed the key role of its Cu^2+^ in mediating this photoresponsive antimicrobial potentiation. Exclusion analysis confirmed the specificity of Cu^2+^ coordination, as other mineral-derived cations (K^+^, Zn^2+^, Mg^2+^, Ca^2+^ and Fe^2+/3+^) and non-metal inorganic constituents (S, B, Si and As) failed to induce significant recovery of activity.

As an essential trace element, Cu^2+^ participates in enzymatic activation and metalloprotein assembly, highlighting the pharmaceutical utility of Cu^2+^ salts [20–24]. Mineral medicinal materials, such as chalcanthite and azurite (Cu_3_(CO_3_)_2_(OH)_2_), demonstrate clinically validated emetic properties, topical antimicrobial efficacy, and wound debridement functions through external application or controlled oral administration [24–26]. Copper gluconate (Cu(C_6_H_11_O_7_)_2_), owing to its high bioavailability, low toxicity, and superior stability, is extensively used in managing copper deficiency syndromes and accelerating tissue regeneration [27, 28]. Transition metal ions (Fe^2+^/Cu^2+^/Mn^2+^/Zn^2+^) form stable coordination complexes with traditional tetracyclines, fundamentally altering their photolysis pathways [11, 29–31]. These metal-ion modified photolytic byproducts exhibit markedly reduced ecotoxicity toward aquatic organisms (e.g., algae, shrimps and fish) compared to parent tetracyclines [30, 31]. This phenomenon in wastewater treatment and soil bioremediation systems provides a chemical basis for developing metal-enhanced tetracycline remediation strategies.

Additionally, milligram-scale Cu^2+^ exhibits broad-spectrum biocidal activity (including bacteria, fungi, protozoa, zooplankton and algae), supporting the use of cupric sulfate in medical device sterilization, disease prevention in aquaculture, and prevention and control of plant diseases and environmental disinfection [32–34]. However, subinhibitory (microgram-scale) Cu^2+^ does not significantly affect bacterial growth. Cu^2+^ may perturb bacterial membrane transport function—a phenomenon whose mechanism needs to be further investigated.

## 2. Results

### 2.1 Exposure to light affects the antibacterial activity of tigecycline

Previous studies have confirmed that tetracycline antibiotics are susceptible to photodegradation and associated activity loss. However, variations in antibacterial activity among different tetracyclines under daylight or artificial lighting have not been reported. In this study, we first evaluated light-induced changes in the antimicrobial activity of five commonly used antibiotics via MIC assays. Notably, only tigecycline exhibited significant differences (*p* < 0.01) in antibacterial activity between light-exposed and dark conditions against the drug-resistant bacteria *Staphylococcus aureus* (*S. aureus*) USA300 and *Escherichia coli* (*E. coli*) ZJ487 (Figure 1A, B). A subsequent test of nine tetracyclines revealed significant antimicrobial activity variations only for minocycline, with other tetracyclines showing no significant light-dependent effects (*p >* 0.05) (Figure 1C and Figure S1). To confirm this observation, pre-illuminated and light-protected versus freshly prepared tigecycline solutions were comparatively tested for 24 h, and the results showed that photolysis is a key factor contributing to the reduced antibacterial efficacy of tigecycline (Figure 1D, E). Thus, developing potential agents that prevent or regulate the photolysis of tigecycline may yield an efficacy equivalent to that of light-protected administration, thereby increasing its antibacterial activity under light exposure.

**Figure 1.**
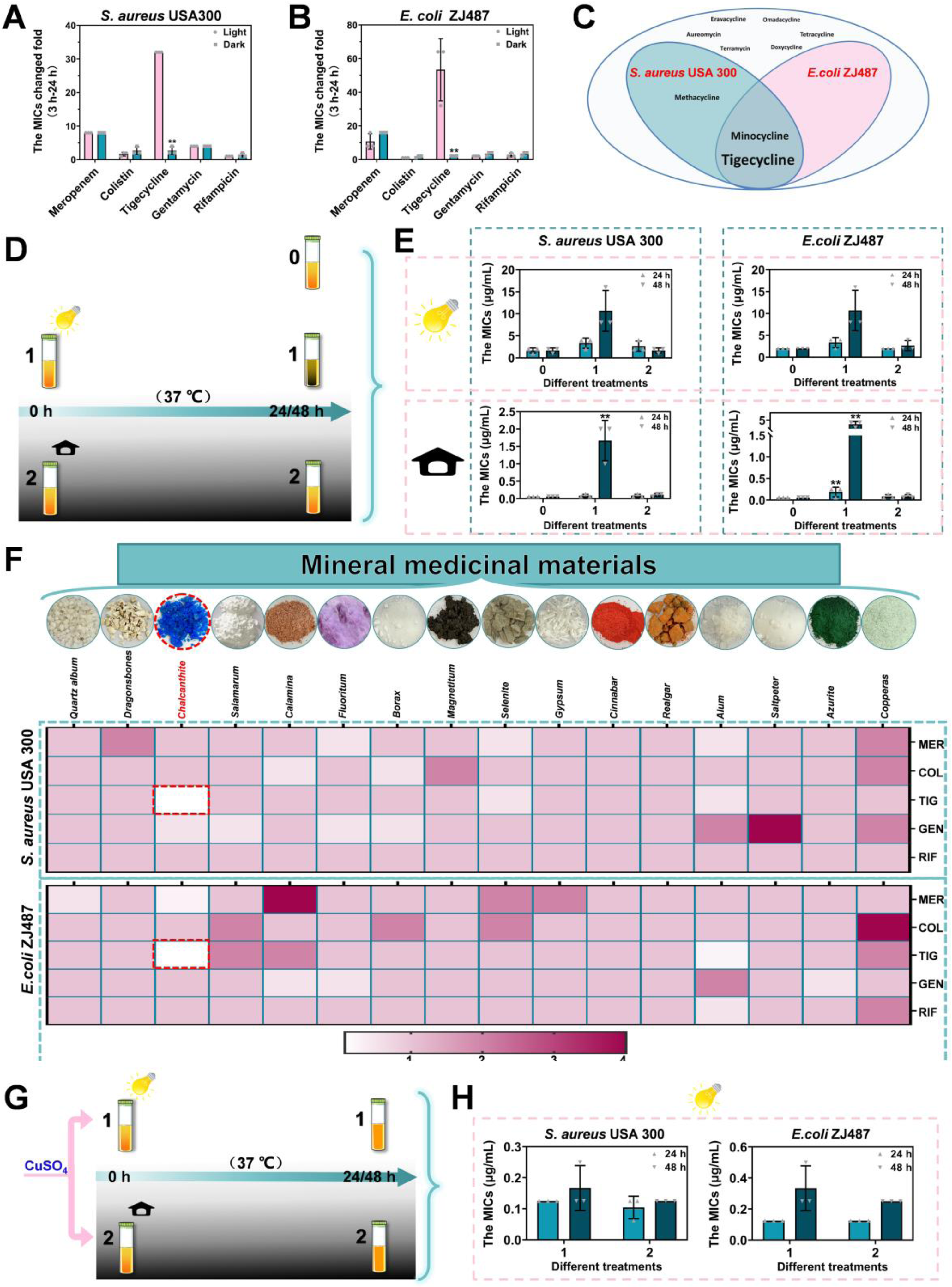
Obtaining bioactive constituents from mineral medicinal materials to counteract tigecycline photolytic inactivation. (**A, B**) The multiple of the increase in the MIC values of meropenem, colistin, tigecycline, gentamycin and rifampicin from 3 h to 24 h under light and dark conditions. (**C**) The Venn diagram illustrates the influence of light on antibacterial efficacy across tetracycline antibiotics (24 h observation). And the larger the text, the greater the influence. (**D, E**) The MICs of tigecycline preparations (0: freshly prepared, 1: light-exposed, 2: light-shielded; Light-exposed and light-shielded for 24 or 48 h) against *S. aureus* USA300 and *E. coli* ZJ487 were quantified under both light and dark conditions, with assessments conducted at 24 h. (**F**) Heat map of FIC index, under light conditions, screening of combinations with broad-spectrum synergistic antibacterial effects among 16 mineral medicinal materials and five antibiotics (MER, Meropenem; COL, Colistin; TIG, Tigecycline; GEN, Gentamycin; RIF, Rifampicin) were screened, and chalcanthite was obtained. (**G, H**) The MICs of tigecycline preparations (all added CuSO_4_, 1: light-exposed, 2: light-shielded, 24 h) against *S. aureus* USA300 and *E. coli* ZJ487 were quantified under light conditions, with assessments conducted at 24 h. Experiments in **A**, **B**, **E**, **F**, and **H** were performed as three biologically independent experiments, and the mean±SD is shown; *P* < 0.05, **, was considered statistically significant.

### 2.2 Medicinal Cu^2+^ salts significantly reverse the antibacterial activity of tigecycline under light conditions

To address the photolysis challenges experienced by tigecycline and minocycline, we performed synergistic antibacterial screening across 16 mineral medicinal materials. As shown in Figure 1F, under controlled photoirradiation, checkerboard MIC testing with five representative antibiotics revealed that exclusively chalcanthite and tigecycline demonstrated considerably higher synergistic efficacy against multidrug-resistant *S. aureus* USA300 and *E. coli* ZJ487 [35, 36]. Subsequent photoirradiation and dark incubation experiments with Cu^2+^ sulfate-supplemented tigecycline, as anticipated, substantially restored the light-compromised antimicrobial activity (Figure 1G, H).

We also systematically investigated the synergistic antimicrobial effects of various Cu^2+^ salts combined with different tetracycline antibiotics against multiple representative pathogenic strains (Figure 2A-F). As shown in Figure 2A, except for *P. aeruginosa* (which possesses specific metabolic mechanisms for cupric ions), Cu^2+^ significantly enhanced the bacteriostatic activity of tigecycline against eight common gram-negative and gram-positive bacteria (number of strains were 23) under light conditions [37, 38]. Concurrently, six commonly used Cu^2+^ salts showed synergistic bacteriostatic effects with tigecycline (Figure 2B), with copper sulfate and copper gluconate exhibiting particular potential pharmaceutical value.

**Figure 2.**
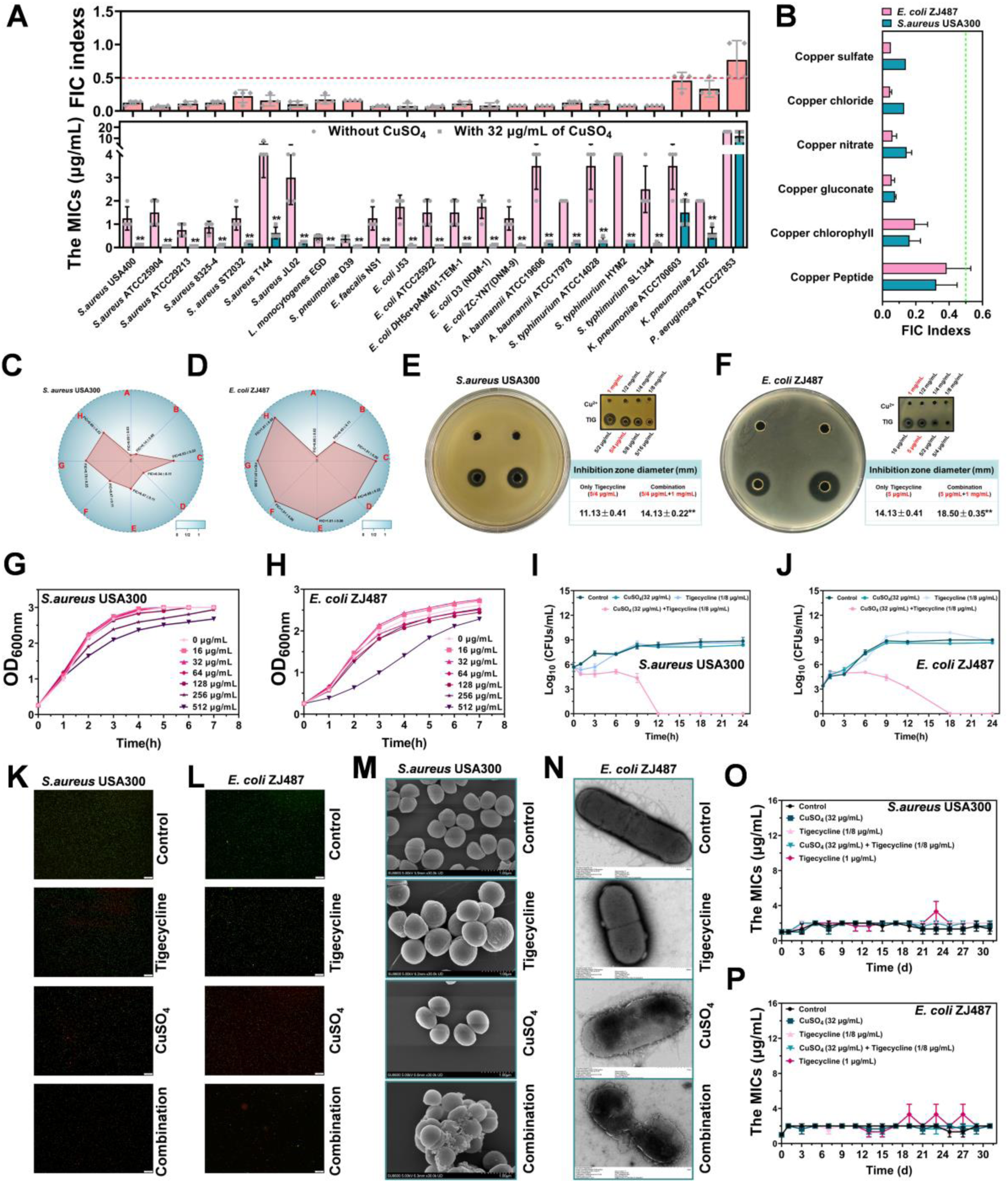
*In vitro* synergistic antibacterial activity of tigecycline combined with Cu^2+^ against multidrug-resistant bacterial strains under light conditions. **(A)** The MIC values and FIC indices of tigecycline with or without 32 μg/mL CuSO_4_ against 23 different strains under light conditions. An FIC < 0.5 is considered to have a synergistic effect. (**B**) The FIC indices of tigecycline with different Cu^2+^ concentrations against *S. aureus* USA300 and *E. coli* ZJ487 under light conditions. (**C, D**) The contribution capacity chart of FIC shows that CuSO_4_ only had an excellent synergistic effect with tigecycline and minocycline against *S. aureus* USA300 and *E. coli* ZJ487 (A, tigecycline; B, minocycline; C, eravacycline; D, aureomycin; E, methacycline; F, terramycin; G, doxycycline; H, tetracycline. n=4). (**E, F**) Optimized agar diffusion assay for tigecycline in combination with CuSO_4_ against *S. aureus* USA300 and *E. coli* ZJ487. (**G, H**) Growth curves for *S. aureus* USA300 and *E. coli* ZJ487. (**I, J**) Time-killing curves for CuSO_4_, tigecycline, combination and control treatment against *S. aureus* USA300 and *E. coli* ZJ487. Fluorescence labeling analysis for *S. aureus* USA300 (**K**) and *E. coli* ZJ487 (**L**) were treated with CuSO_4_ (32 µg/mL), tigecycline (1/8 µg/mL), the combination or control (bacteria without any treatment), live bacteria were dyed green and dead bacteria were dyed red (scale bar = 50 μm). (**M, N**) SEM (scale bar = 1.00 μm) and TEM (scale bar = 1.00/0.50 μm) analyses of *S. aureus* USA300 and *E. coli* ZJ487 were also performed with the above treatments. (**O, P**) CuSO_4_ (32 µg/mL), tigecycline (1/8 µg/mL), the combination (32 µg/mL+1/8 µg/mL), or tigecycline (1 µg/mL) did not induce the resistance of bacteria to tigecycline under light conditions; Experiments were performed as three or four biologically independent experiments, and the mean±SD is shown; * indicates *P* < 0.05. ** indicates *P* < 0.01.

As the antibacterial activity of conventional tetracyclines other than tigecycline and minocycline did not significantly differ between photic and aphotic conditions, these compounds exhibited lower or no synergistic effects on *S. aureus* USA300 and *E. coli* ZJ487 in the presence of Cu^2+^ (Figure 2C, D). Other tetracyclines lacked synergy with Cu^2+^, possibly due to the absence of a dimethylamino group on the D-ring structure (Figure 2C, D). The enhanced bacteriostatic efficacy of Cu^2+^ on tigecycline under light conditions was further validated through disk diffusion antimicrobial susceptibility testing (Figure 2E, F). The synergistic bactericidal effect of Cu^2+^ with tigecycline against *S. aureus* USA300 and *E. coli* ZJ487 was investigated through time-killing assays, live/dead bacterial staining, scanning electron microscopy (SEM), and transmission electron microscopy (TEM). The results of the growth curve indicated that CuSO_4_ at concentrations of ≤256 μg/mL did not significantly affect the growth of the tested bacteria (Figure 2G, H). Photic exposure revealed that neither Cu^2+^ nor tigecycline alone induced effective bacterial killing in the culture media, whereas their combination achieved significant bactericidal efficacy (Figure 2I-L). Electron microscopy further demonstrated that bacterial aggregation, morphological aberrations, membrane rupture, and cell death occurred in the combination group (Figure 2M, N). Serial passage experiments over 30 days confirmed that neither tigecycline, Cu^2+^ alone, nor their combination under light conditions induced bacterial resistance to tigecycline (Figure 2O, P).

Additionally, Tet(X3/X4)-producing bacteria, which confer high-level resistance to tigecycline through enzymatic inactivation mechanisms, and bacteria extremely sensitive to tigecycline even under photoirradiation, such as *Streptococcus mutans* (*S. mutans*) (a facultative anaerobe whose increased susceptibility to Cu^2+^-mediated bactericidal effects under oxygen-deprived conditions), demonstrated negligible photolysis of tigecycline (Figure S2A-F). And other metal ions that complex with tetracyclines do not have effective synergistic effects (Figure S2G).

### 2.3 The regulatory effect of Cu^2+^ on tigecycline facilitates accelerated photolysis while concurrently generating products with antibacterial activity degradation ability and antiphotolytic properties

We systematically investigated the Cu^2+^-mediated photochemical dynamics of tigecycline through spectroscopy and chromatography analysis. Cu^2+^ addition induced a pronounced chromatic darkening of the tigecycline solution after 24 h under light conditions (Figure 3A). And Cu^2+^ supplementation induced characteristic hyperchromic effects (260-280 nm and 345-375 nm peak intensification) with bathochromic shifts, which indicated metal-ligand coordination complex formation between tetracyclines and Cu^2+^ before the reaction (Figure 3B, C). After 24 h of photoirradiation, Cu^2+^-tigecycline/doxycycline systems exhibited attenuation at 260-280 nm and 345-375 nm (dashed-line circles) concomitant with emerging absorption shoulders at 530 nm (black arrows), suggesting that Cu^2+^-mediated reorganization of π→π* electronic transitions and potential n→π* charge transfer states occurred during photolytic pathway modification (Figure 3B, C). The high-performance liquid chromatography (HPLC) results showed that,compared to control conditions involving only photoirradiation or Mg^2+^ supplementation, the introduction of Cu^2+^ after 24 h of photoexposure significantly increased the photolysis of tigecycline (Figure 3E). This Cu^2+^-facilitated photolysis process generated distinct photolysis products, as indicated by UV-Vis spectral analysis (Figure 3E). Mass spectrometry detection confirmed that the Cu^2+^ supplementation substantially promoted the generation of photolysis byproducts (Figure 3F, G).

**Figure 3.**
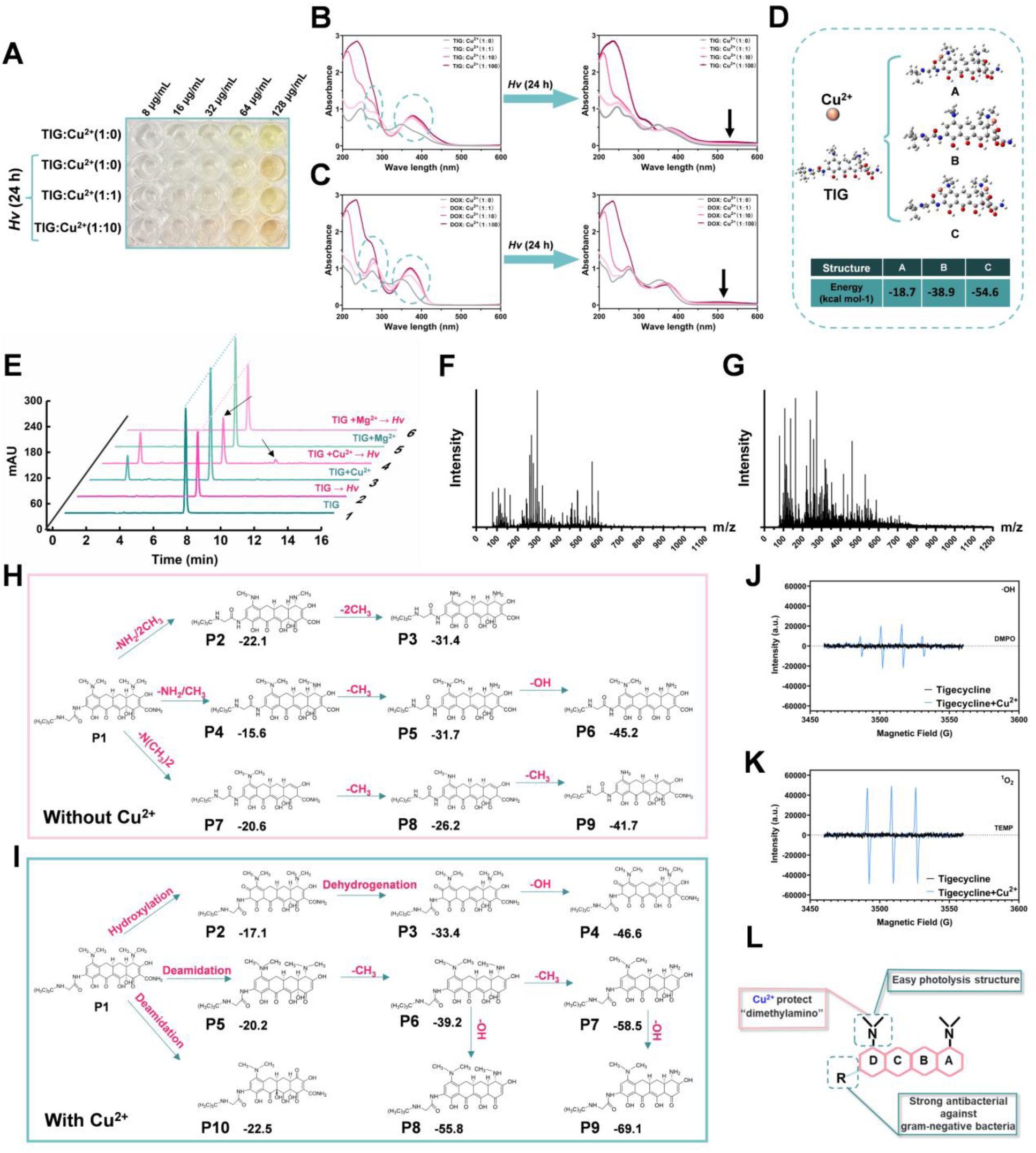
The Cu^2+^-mediated acceleration of tigecycline photolysis. (**A**) Representative color of tigecycline with or without CuSO_4_ under light conditions for 24 h. (**B, C**) UV–visible absorption spectra (200-600 nm) of different tigecycline/doxycycline -CuSO_4_ ratios, fresh or exposed under light conditions for 24 h. (**D**) Potential binding conformations of copper ions with tigecycline and computational quantification of adsorption energies. (**E**) The HPLC results of tigecycline after different treatments are presented (MgSO_4_ was used as a control). High-resolution mass spectrometric characterization of tigecycline with (**G**) or without (**F**) CuSO_4_ after 24 h of photoirradiation. Photolysis products and degradation pathways of tigecycline with (**I**) or without (**H**) CuSO_4_ under light irradiation. Electron paramagnetic resonance (EPR) characterization of hydroxyl radicals (·OH) (**J**) and singlet oxygen (^1^O_2_) (**K**) was performed using the spin-trapping agents DMPO and TEMP across experimental groups after treatment for 30 min under light conditions. (**L**) Schematic representation: Elucidating the Cu^2+^-induced protection mechanism of the bioactive scaffold of tigecycline combined with antibacterial efficacy restoration, supplemented by molecular mapping of essential antigram-negative functionalities.

We applied the application of quantum chemical computations to elucidate the molecular interactions between tigecycline and Cu^2+^ and performed the structural characterization of photolysis byproducts retaining antibacterial efficacy (Figure 3H, I). The divergent photolytic pathways under Cu^2+^-supplemented versus Cu^2+^-free conditions, revealed a protective coordination mechanism where Cu^2+^ preferentially stabilizes the dimethylamino moieties (strong adsorption energy) (Figure 3D, H, I). This stabilization effect aligns with our initial hypotheses. The absence of D-ring dimethylamine groups in doxycycline fundamentally negates the capacity of Cu^2+^ to preserve antibacterial activity during photoirradiation (Figure S3). By analyzing the mass spectrometric data, we qualitatively identified 10 Cu^2+^-associated versus nine Cu^2+^-free photodegradation products, validating the computational pathway predictions (Figure S4, S5). Additionally, photoirradiation processes generate reactive oxygen species (ROS) predominantly comprising singlet oxygen (^1^O_2_) and hydroxyl radicals (·OH), which subsequently elicit oxidative cascade reactions through electron transfer mechanisms. Electron paramagnetic resonance (EPR) spin-trapping quantification revealed greater yields of both ^1^O_2_ and ·OH under Cu^2+^-supplemented conditions, indicating that Cu^2+^ induced more ^1^O_2_ and ·OH production (Figure 3J, K; Figure S2H, S2I). This ensemble of experimental evidence mechanistically establishes that inherent photolysis reduces the antibacterial activity of tigecycline and minocycline in solution-phase systems because of the structural vulnerability of their D-ring dimethylamino moieties, with degradation pathways directly correlated with the loss of antimicrobial efficacy. Crucially, Cu^2+^ effectively mitigates photoinduced structural rearrangement through chelation-induced frontier orbital modulation, thereby preserving the pharmacophoric integrity necessary for antibacterial activity (Figure 3L).

### 2.4 Confirmation of the effect of Cu^2+^ on the formation of bacterial biofilms

Initially, we conducted employed crystal violet staining and fluorescence staining assays to demonstrate that Cu^2+^ alone or in combination with tigecycline effectively inhibited the formation of *S. aureus* USA300 and *E. coli* ZJ487 biofilms under light irradiation (Figure 4A, B). Moreover, quantitative analysis of the biofilm content and bacterial load in the biofilm confirmed the antibiofilm activity of Cu^2+^, either alone or synergistically with tigecycline (Figure 4C-F). The leakage of bacterial DNA and proteins following treatment with Cu^2+^ or tigecycline revealed that their combination significantly disrupted bacterial membrane integrity, thereby causing extracellular release of DNA and proteins (Figure 4G, H). Notably, Cu^2+^ exhibited potent disruptive activity against *S. aureus* USA300 biofilms even at non-antibacterial concentrations (Figure 4G, H).

**Figure 4.**
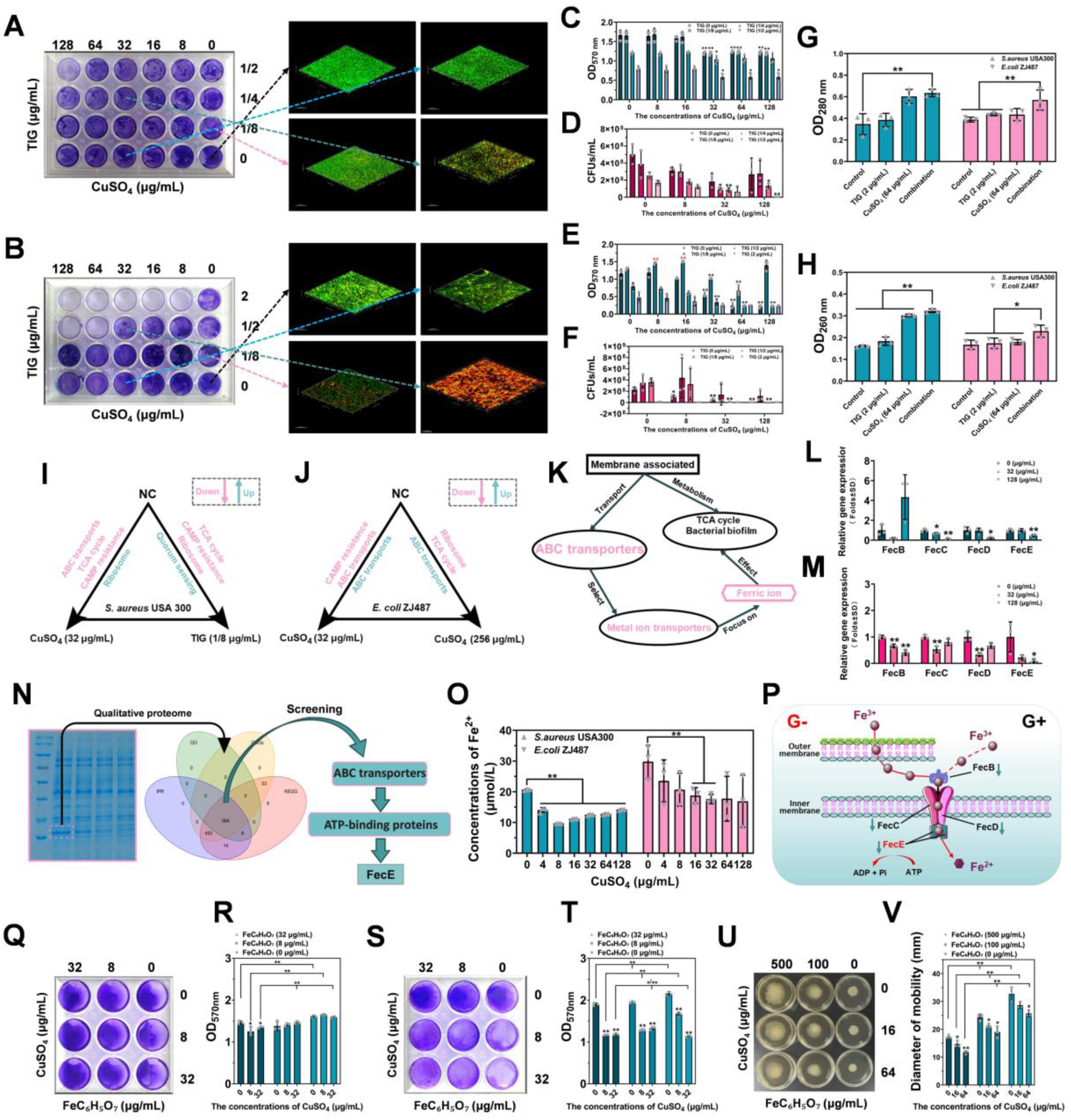
Effects of Cu^2+^ alone, or in combination with tigecycline on biofilm formation, and on the functional integrity of the bacterial cellular cell membrane. Crystal violet (CV)-stained biofilms and confocal microscopy images of *S. aureus* USA300 (**A**) and *E. coli* ZJ487 (**B**) biofilms grown with different concentrations of CuSO_4_ and/or tigecycline under light conditions. The fluorescence intensity of each sample represents the thickness of the biofilm. The graphs present the OD_570_ values of crystal violet (CV)-stained biofilms (**C, E**) and biomass (CFUs) quantified by microbiological plating (**D, F**). Measurement of cellular leakage of protein (**G**) and nucleic acid (**H**) from *S. aureus* USA300 and *E. coli* ZJ487 treated with different concentrations of CuSO_4_ and/or tigecycline under light conditions. Transcriptomic profiling of the tested bacteria following exposure to CuSO_4_ (32 µg/mL) and the common downregulated pathways from *S. aureus* USA300 (**I**) and *E. coli* ZJ487 (**J**) were obtained. Untreated and CuSO_4_-treated (256 µg/mL) or tigecycline-treated (1/8 µg/mL) samples were used as controls. (**K**) The ABC transporter associated with bacterial energy metabolism, pathogenicity, and biofilm formation was targeted, and the ferric citrate transporter was found to be downregulated in *S. aureus* USA300 and *E. coli* ZJ487. RT-qPCR was performed to analyze the effects of different concentrations of CuSO_4_ on the expression of key proteins in FecABCDE from *S. aureus* USA300 (**L**) and *E. coli* ZJ487 (**M**). (**N**) Following precise excision of semiquantitatively differential protein gel bands, in-depth proteomic profiling was performed, integrated with multilayered bioinformatic interrogation, and the *FecE* gene was identified in the ferric citrate transporter assembly. (**O**) Determination of ferrous ion content in the tested bacteria. (**P**) Cu^2+^ can regulate multiple genes affecting Fe^3+^ transport. After adding ferric citrate and CuSO_4_, the biofilm content of *S. aureus* USA300 (**Q, R**) and *E. coli* ZJ487 (**S, T**) was detected via CV staining, and the effects on the surface motility of *A. baumannii* ATCC17978 (**U, V**) were also detected simultaneously. Experiments dates were shown as mean±SD. * indicates *P* < 0.05. ** indicates *P* < 0.01.

### 2.5 Elucidating the mechanisms underlying Cu^2+^-mediated modulation of bacterial iron ion transport homeostasis

Non-antibacterial concentrations of Cu^2+^ perturb bacterial biofilm formation, probably by modulating efflux systems or membrane transporters. We systematically investigated the transcriptomic profiles of *S. aureus* USA300 and *E. coli* ZJ487 exposed to different concentrations of Cu^2+^ (high/low concentrations) and sub-MICs of tigecycline. Comparative analysis revealed significant downregulation of two biofilm-associated regulatory systems in Cu^2+^-treated *S. aureus* USA300 and *E. coli* ZJ487: ABC transporters and cationic antimicrobial peptide (CAMP) resistance pathways (Figure 4I, J; Figure S6). Given the cationic nature of Cu^2+^, we focused on deciphering the regulatory effect of copper on the ABC transporter. Further mining of omics data revealed Cu^2+^-induced transcriptional perturbations in multiple metal ion transporter genes, particularly within the ferric citrate transport system (Figure 4K). Subsequent RT-qPCR assays targeting four genes (*fecB*, *fecC*, *fecD*, and *fecE*) in fecBCDE-mediated iron transport in both *S. aureus* USA300 and *E. coli* ZJ487 demonstrated suppression by low-concentration Cu^2+^ (Figure 4L, M).

SDS-PAGE analysis of the Cu^2+^-treated *E. coli* ZJ487 cultures revealed that the protein band intensity decreased considerably in a concentration-dependent manner (Figure S7A-S7C). Targeted proteomic characterization of this differential band through multi-strategy analyses identified FecE, an ATP-binding protein of the fecBCDE transporter, as the primary constituent (Figure 4N; Figure S8). Quantitative analysis of ferrous ion levels demonstrated that the presence of Cu^2+^ ions at non-bacteriostatic concentrations significantly decreased the intracellular ferrous ion content in bacterial cells (Figure 4O). These findings collectively demonstrate that copper ions regulate the expression of genes associated with bacterial iron transporters (represented by FecBCDE), thereby suppressing iron acquisition and utilization in bacterial cells (Figure 4P).

Iron deprivation has multiple physiological consequences as iron plays a crucial role in maintaining bacterial membrane integrity, energy transduction, and the deployment of virulence factors. Exogenous ferric citrate supplementation reversed these phenotypes, increasing the biofilm biomass of *S. aureus* USA300 and *E. coli* ZJ487, while increasing the surface motility of *Acinetobacter baumannii* (*A. baumannii*) ATCC17978 (Figure 4Q-V). Conversely, had diametrically opposite effects on the integrity of the outer membrane of *E. coli* ZJ487 and reduced the level of intracellular ATP; these effects were not detected in *S. aureus* USA300 due to its lack of an outer membrane permeability barrier (Figure 4Q-V; Figure S7D-S7G). Divalent cations (Mg^2+^ and Zn^2+^) failed to suppress the surface-associated motility of *A. baumannii* ATCC17978 (Figure S7H-S7K), and even promoted sliding (Ca^2+^) (Figure S7L, S7M).

### 2.6 Cytotoxicity of Cu^2+^ and tigecycline and their combined cytoprotective efficacy at the cellular level

We initially evaluated the cytotoxicity of tigecycline and pharmaceutical-grade Cu^2+^ salts to A549 cells under various experimental conditions. The results revealed demonstrated no cytotoxicity toward the tested cell within 6 h when tigecycline was ≤ 2 μg/mL and copper gluconate was ≤ 64 μg/mL (Figure S9A, S9B). Light exposure induced a mild increase in the cytotoxic potential of tigecycline, whereas copper gluconate administration partially attenuated this increase in light-induced toxicity. However, neither phenomenon was statistically significant (*p*>0.05). Furthermore, tigecycline in combination with copper gluconate significantly inhibited the cellular invasion of RAW264.7 cells by both *S. aureus* USA300 and *E. coli* ZJ487 at critical timepoints (2, 6, and 12 h post-treatment) (Figure S9C, S9D). Collectively, these findings confirmed that co-administration of tigecycline with medicinal Cu^2+^ formulations at non-toxic concentrations effectively preserves cellular integrity to prevent bacterial pathogenesis.

### 2.7 Efficacy of copper gluconate combined with tigecycline *in vivo*

To determine the substantial synergistic antimicrobial efficacy of copper ions combined with tigecycline under light conditions *in vitro*, we subsequently investigated their therapeutic potential *in vivo*. We established two distinct infection animal models (Figure 5A, B), with all experimental procedures conducted under continuous light exposure. Copper gluconate at ≤ 50-fold therapeutic concentrations elicited no observable toxicity in *Galleria mellonella* (*G. mellonella*) larvae (Figure S9E-S9G). An appropriate lethal dose of *E. coli* ZJ487 was administered to *G. mellonella* larvae, followed by the initiation of various treatments 2 h post-infection (Figure 5A; Figure S9H). The larvae treated with copper gluconate and tigecycline at 1:1, 1:5, and 1:10 molar ratios presented survival rates that increased from 35% to ≥80% at 7 days after treatment (Figure 5C). Simultaneously, quantitative bacteriological analysis revealed significant attenuation of bacterial loads in the combined therapy group compared to the monotherapy or control groups (Figure 5D).

**Figure 5.**
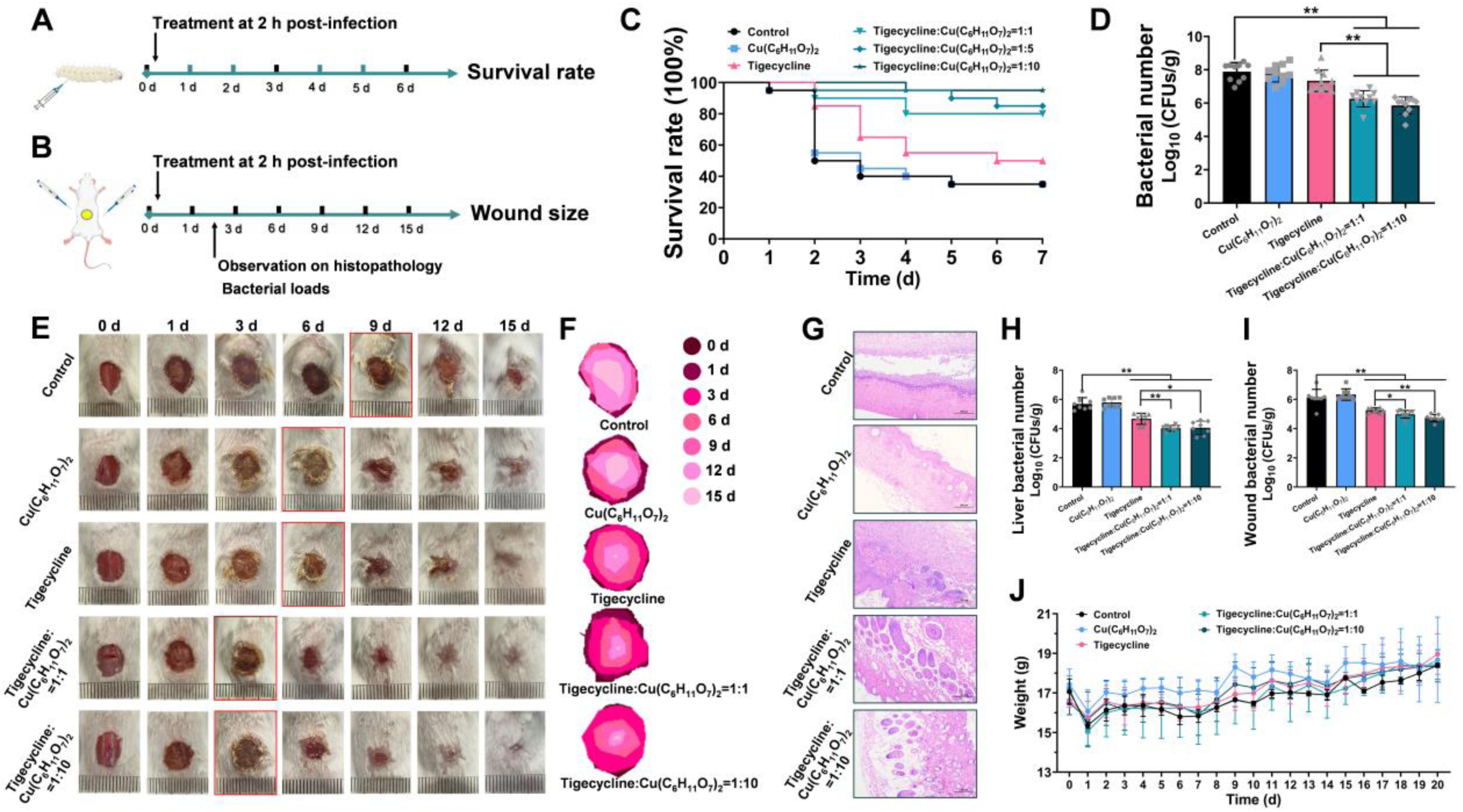
Copper gluconate restores the antimicrobial activity of tigecycline against different drug-resistant pathogen infections under light conditions. Scheme of the experimental protocol for the *G. mellonella* larvae infection model (**A**) and mouse skin wound infection model (**B**). (**C**) Survival curves of *G. mellonella* larvae during 7 days after infection (n = 20). (**D**) Bacterial loads in *G. mellonella* larvae at 48 h post-infection (n = 10). (**E)** Representative images of wound healing recorded 15 days after infection. (**F**) Traces of wound closure in *S. aureus* USA300 infected mice after treatment. (**G**) Representative images of H&E-stained of the wound tissue used to measure epithelialization. Bacterial loads in livers (**H**) and wounds (**I**) after 48 h of treatment. (**J**) The body weight of the mice was monitored over 20 days (n = 3, Stochastic). * indicates.

In the mouse wound infection model, the copper-tigecycline combination demonstrated better therapeutic outcomes in terms of multiple parameters: enhanced wound morphology restoration, reduced lesion dimensions, and mitigated histopathological damage relative to those of the monotherapy groups (Figure 5E-G). The combinations (1:1 and 1:10) also significantly decreased the wound bioburden (*p* < 0.05) (Figure 5H, I). No significant intergroup differences in body weight trajectories were observed throughout the 20-day monitoring period (Figure 5J). These findings confirmed the therapeutic potential of Cu^2+^ in enhancing the antibacterial activity of tigecycline under light conditions, thereby providing a promising strategy for combating clinical complex drug-resistant bacteria *in vivo* (Figure 6).

**Figure 6.**
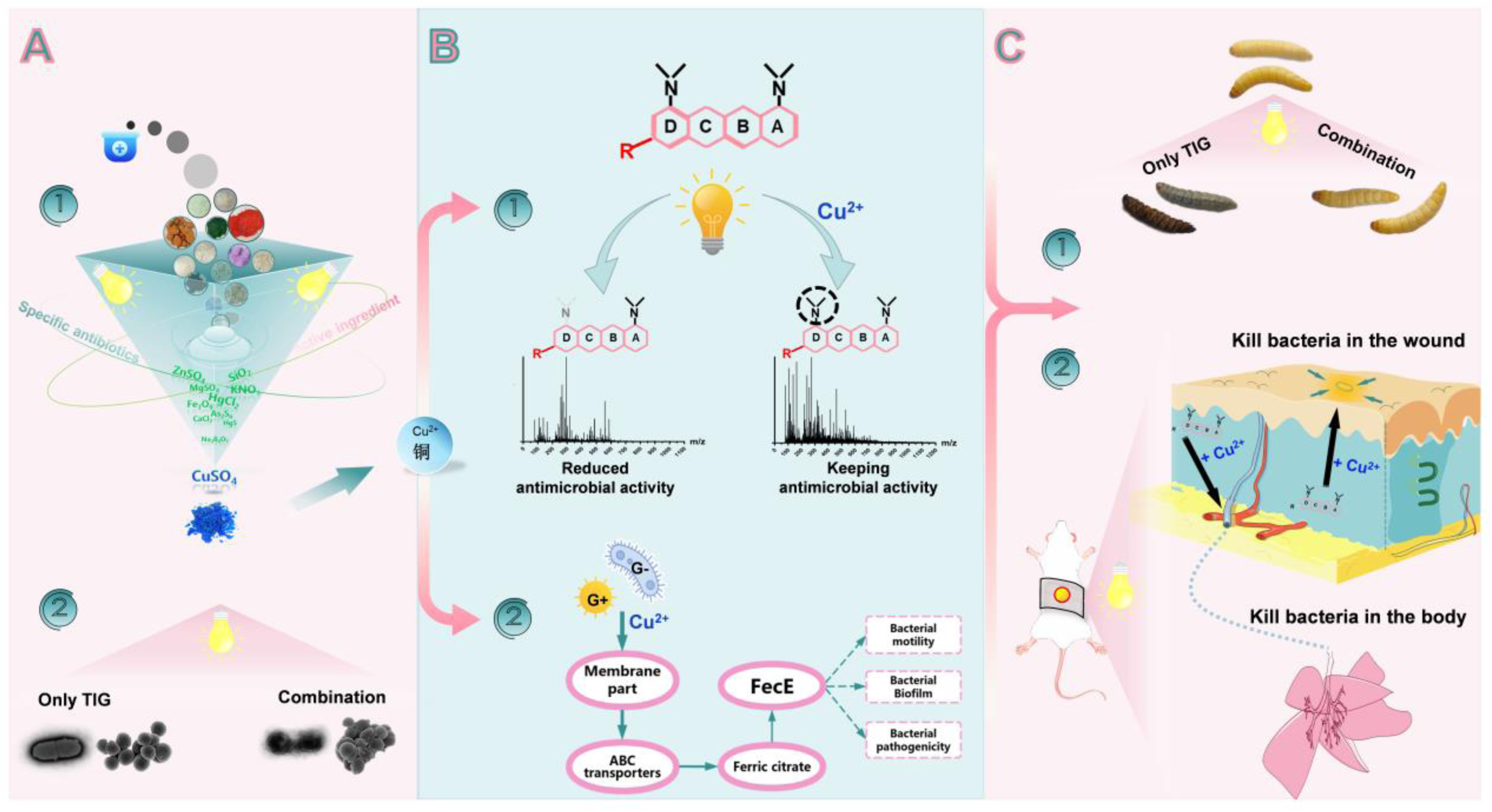
Schematic model of dual-mechanism potentiation: Cu^2+^-mediated enhancement of tigecycline bactericidal activity under photodynamic conditions.

## 3. Discussion

A seminal publication cautions that antimicrobial resistance (AMR) may directly contribute to more than 39 million global fatalities annually by 2050 [39]. This therapeutic limitation highlights an urgent imperative to develop mechanism-based antimicrobial agents through novel therapeutic targets or optimize existing antibiotic administration protocols through enhanced management frameworks [39–42]. Tigecycline is a structural analog within the tetracycline class and is universally recognized as a first-line broad-spectrum antimicrobial agent in clinical practice because of its cost-effectiveness and therapeutic efficacy [43, 44]. However, photolytic decomposition is considerably accelerated under prolonged exposure to different types of light (sunlight, artificial light) and atmospheric oxygen. Such molecular transformations not only compromise the intrinsic antimicrobial potency but also introduce substantial variability in the evaluation of antibacterial efficacy, thereby necessitating stringent storage protocols to preserve the integrity of compounds. Therefore, overcoming the inherent photolytic vulnerability of tigecycline pharmacophores may lead to the de novo development of a mechanistically distinct broad-spectrum antimicrobial agent with greater enhanced physicochemical stability.

The dimethylamino group at the C-7 position of the D-ring in minocycline is structurally so advantageous that this moiety has been retained or modification in nearly all subsequent semisynthetic and fully synthetic tetracycline derivatives [19, 45, 46]. The topical ointment formulation of minocycline has been approved for periodontitis management [47, 48]. The resolution of photolytic instability in minocycline scaffolds not only preserves their antimicrobial pharmacophores but also unlocks formulation engineering possibilities. Notably, p-toluenesulfonic acid (TsOH), a non-oxidizing organic strong acid, plays dual pharmaceutical roles as a stabilizer and a reaction intermediate. Its incorporation into the molecular architecture of omadacycline maintains the structural integrity of omadacycline, effectively reducing photosensitivity. In contrast, eravacycline represents an alternative stabilization strategy via structural substitution of the dimethylamino group with fluoride anions at C-7, achieving comparable photostability.

Cu^2+^ salt, such as copper gluconate, copper sulfate, copper chloride, and copper chloride, have been extensively used as feed additives to increase feed conversion efficiency and promote organismal growth and development. The Joint FAO/WHO Expert Committee on Food Additives (JECFA), and the National Food Safety Standard of China has endorsed copper gluconate as a food nutritional enhancer characterized by greater utility, safety, and essentiality, justifying its selection as the copper source in our animal experiments (GB 1903.8-2015, China) [28]. Those with high therapeutic value do not include Cu^2+^-peptide complexes. As the glycyl-histidyl-lysine (GHK) peptide has a high affinity for divalent Cu^2+^, it spontaneously coordinates with Cu^2+^ to form the GHK-Cu^2+^ complex. This interaction constrains the photodegradation regulatory effect of free Cu^2+^ on the potentiation of the antibacterial activity of tigecycline. Our investigation revealed a non-linear dose-response relationship for copper ion supplementation, with checkerboard MIC assays demonstrating paradoxical attenuation of synergistic efficacy at concentrations ≥256 μg/mL. This inverse pharmacodynamic profile beyond the threshold concentration suggests the activation of multifactorial counter-regulatory processes, possibly involving microbial stress-response pathways or ligand-saturation effects. The observed biphasic behavior highlights the necessity for precise optimization of copper dosing parameters in antimicrobial combinatorial therapeutic techniques.

The Fe^2+^/Fe^3+^ ions are critical determinants of bacterial stress adaptation, biofilm formation, and virulence expression [49, 50]. They operate through tightly regulated acquisition systems [50]. The Cu^2+^-induced dysregulation of the Fe^3+^ acquisition transporter (e.g., FecABCDE) prompted the hypothesis that Cu^2+^ disrupts iron homeostasis [51, 52]. Notably, nonferrous metal ions (Mg^2+^, Zn^2+^, and Ca^2+^) showed negligible iron homeostasis interference in parallel assays, highlighting the specificity of Cu^2+^ metalloregulatory crosstalk (Figure S2G; Figure S7H-S7M). Additionally, proteomic profiling revealed significant enrichment of ABC transporter-associated proteins, including FecE, LptB, HisP, GlnQ, DsdX, PhoU, YbbA, and ArtP. This observation suggests that Cu^2+^ may exert broad-spectrum regulatory effects on bacterial energy metabolism and membrane transport processes, whereas our investigation characterized a singular mechanistic interaction involving the ferric citrate transport machinery.

## 4. Conclusion

Except for the strains of *P. aeruginosa*, Tet(Xs)-positive bacteria, and extremely sensitive tetracycline bacteria, our results provided evidence for advancing the development of topical tigecycline formulations while delineating pharmaceutically optimized Cu^2+^ complexes as strategically viable candidates for designing antibacterial adjuvants. The demonstrated capacity of Cu^2+^-based methods to enhance the antimicrobial activity of tigecycline under light conditions highlights their potential as therapeutic enhancers in photoresponsive antimicrobial regimens, offering a paradigm shift in the optimization of combination therapy against multidrug-resistant pathogens.

## 5. Experimental Section

### 5.1 Bacterial strains, chemicals and reagents

The origins of all bacterial strains used in this study are detailed in Table S1. Gram-negative bacterial strains including *E. coli, A. baumannii*, *S. typhimurium*, *K. pneumoniae* and *P. aeruginosa* were grown under agitation in LB medium (Sigma-Aldrich, USA) at 37 °C. Gram-positive bacterial strains, including *S. aureus*, *L. monocytogenes*, *S. pneumoniae*, and *E. faecalis*, were grown in TSB medium (Qingdao Hopebio Co., Ltd., China). Additionally, 5% serum was added, when necessary, at 37 °C. *Streptococcus mutans* was grown in an anaerobic environment in BHI medium (Qingdao Hopebio Co., Ltd., China) at 37 °C.

All the antibiotics were purchased from the National Institutes for Food and Drug Control (China), Shanghai Yuanye Biotechnology Co., Ltd. (China), and Dalian Meilun Biotechnology Co., Ltd. A ROS assay kit (S0033S) and an ATP assay kit (S0026) were obtained from Beyotime Biotechnology Co., Ltd., China. The Cell Ferrous Iron Colorimetric Assay Kit was purchased from Wuhan Yilerite Biotechnology Co., Ltd., China. Chemical reagents were purchased from Beijing Chemical Works (China), Shanghai Macklin Biochemical Co., Ltd. (China) and Sigma-Aldrich (USA). The mineral-based traditional Chinese medicine used was obtained from the Affiliated Hospital of Nanjing University of Chinese Medicine.

### 5.2 Cell lines and animals

Lung carcinoma cells (A549) (RRID: CVCL_0023) were purchased from the American Type Culture Collection (ATCC, USA) and were cultured using a nutrient-rich mixture of DMEM supplemented with 10% fetal bovine serum (Biological Industries, BI), penicillin (100 U/mL), and streptomycin (100 μg/mL). The cells were cultured at 37 °C in an incubator with a 5% CO_2_ atmosphere under constant humidity.

The *G. mellonella* larvae (450±50 mg) were purchased from Keyun Biology Co., Ltd. (China). All larvae exhibited no observable melanization and demonstrated active motility, with initial mean body weights remaining equivalent across all experimental cohorts. Female BALB/c mice (6-8 weeks old) were purchased from GemPharmatech (Nanjing, China). The mice were acclimated for five days in a standard feeding environment with a settled light-dark cycle (12 h:12 h), at a temperature of 24 ±2 °C and a humidity of 55 ±10%. All animal procedures were conducted following protocols approved by the Ethics Review Committee of Animal Experimentation at Ningxia University (No. NXU-2025-062) and in compliance with the principles of the Animal Welfare Act.

### 5.3 Antimicrobial susceptibility assay

The MICs of the candidate mineral medicinal materials, ingredients, and different antibiotics for the tested bacterial strains were determined by the standard broth microdilution method based on the Clinical and Laboratory Standards Institute (CLSI) guidelines [53]. Briefly, candidate compounds or antibiotics were subjected to twofold serial dilutions in sterile 96-well microtiter plates containing Mueller-Hinton broth (MHB), followed by the addition of 100 μL overnight-cultured bacterial suspensions (1×10^6^ CFUs/mL) to each well. After 16-24 h of static coincubation at 37 °C under photic (fluorescent light: 11 W/m^2^) or aphotic conditions, the lowest concentrations demonstrating no visible bacterial growth were validated as the MICs of the test agents. The synergistic effects of the candidate compounds and antibiotics across all tested strains were systematically assessed by the checkerboard microdilution method. The fractional inhibitory concentration index (FICI) was calculated using the following formula: FIC index = (MIC of candidate compounds in combination/MIC of candidate compounds alone) + (MIC of antibiotics in combination/MIC of antibiotics alone). A synergistic interaction was defined as an FICI value ≤ 0.5. Additionally, the Oxford cup method was also used to confirm the synergistic effect between Cu^2+^ and tigecycline.

### 5.4 Growth curves and resistance development assay

For the growth curve assay, *S. aureus* USA300 and *E. coli* ZJ487 were cultured in the corresponding broth media, and calibrated to an initial inoculum density of 0.3 optical density units at 600 nm. Sterile-filtered CuSO_4_ solutions were supplemented with prepared bacterial suspensions to achieve final concentrations ranging from 16 μg/mL to 512 μg/mL. The bacterial cultures were co-cultured at 37 °C with shaking at 200 rpm. The absorbance of each sample at 600 nm was monitored every 60 min. Growth curves were plotted with triplicate measurements [54].

For serial passage resistance test, *S. aureus* USA300 and *E. coli* ZJ487 were subsequently serially passaged (31 generations) under sub-inhibitory tigecycline (1/8 μg/mL or 1 μg/mL) and/or Cu^2+^ (32 μg/mL) in light environments to model evolutionary adaptation dynamics [36, 55]. Post-evolution MIC was determined under standardized photic conditions using broth microdilution. Resistance trajectory modeling delineated tigecycline and/or Cu^2+^ resistance evolution pathways.

### 5.5 Time-killing assays

The quantifiable bactericidal effect of tigecycline combined with Cu^2+^ was evaluated by conducting time-killing assays [55]. Briefly, *S. aureus* USA300 and *E. coli* ZJ487 in the mid-logarithmic phase were separately diluted to 5 × 105 CFU/mL in 96-well plates in MHB medium, and tigecycline (1/8 μg/mL), CuSO4 (32 μg/mL), tigecycline + CuSO4 (1/8 μg/mL + 32 μg/mL) or a blank control (0 μg/mL + 0 μg/mL) was added. The bacterial cultures in 96-well plates were incubated at 37 °C and removed at 0 h, 1 h, 3 h, 6 h, 9 h, 12 h, 18 h, and 24 h post-inoculation for bacterial counts. The bacterial cultures from different treatment groups were subjected to 10-fold serial dilutions, and 20 μL of each diluted culture was spread onto agar plates. Colony-forming units (CFUs) were quantified after overnight incubation at 37 °C.

### 5.6 Live/dead bacteria staining

The bacteria *S. aureus* USA300 and *E. coli* ZJ487 in the mid-logarithmic phase were treated similarly to the method followed in the time-killing assays described above. The bacteria were collected, resuspended, and stained using the LIVE/DEAD BacLight Bacterial Viability Kit (Invitrogen) following the manufacturer’s instructions. Live bacteria were labeled with green fluorescence, and dead bacteria were labeled with red fluorescence. Fluorescence staining and visualization were also performed on biofilm-associated bacteria to characterize their spatial distribution and ability to survive.

### 5.7 SEM assays and TEM analysis

*S. aureus* USA300 and *E. coli* ZJ487 were treated with tigecycline (1/8 μg/mL), CuSO_4_ (32 μg/mL), combination (1/8 μg/mL + 32 μg/mL), or a blank control (0 μg/mL + 0 μg/mL) in TSB/LB medium for 24 h, then washed with sterile PBS buffer, and resuspended to obtain an OD_600nm_ = 0.5. The samples were incubated with polylysine (0.1%), resuspended in PBS, and fixed in glutaraldehyde overnight at 4 °C. The changes in the external morphology of *S. aureus* USA300 after different treatments were observed via SEM (SU8600, Hitachi Co., Ltd., Japan). For TEM analysis, the bacterial cultures of *E. coli* ZJ487 were fixed with sterile glutaraldehyde phosphate. Each sample (10 μL) was spread on TEM grids and incubated for 10 min at room temperature. Uranyl acetate/lead was used to stain the bacterial samples, and a transmission electron microscope (HT7700, Hitachi Co., Ltd, Japan) was used to observe the internal variation in *E. coli* ZJ487.

### 5.8 Analytical methods

Aqueous solutions containing different concentrations of Cu^2+^, tigecycline, or doxycycline, either individually or in combination, were dissolved in distilled water and exposed to fluorescent light irradiation at 37 °C for 24 h. Freshly prepared, non-irradiated controls corresponding to each treatment were established in parallel. Colorimetric changes across all groups were monitored visually, and UV-Vis absorption spectra were measured using a spectrophotometer (PERSEE, China) over the range of 200-600 nm with a wavelength resolution of 1 nm.

An HPLC system (1260 Infinity II, Agilent, USA) equipped with a C18 column (5 μm, 4.6 × 250 mm) was used to detect and analyze the changes in the content of the photolysis product of tigecycline with/without Cu^2+^ in light or darkness. The concentrations of tigecycline and its photolysis products were determined via HPLC at 248 nm. The mobile phase was acetonitrile and phosphate buffer (0.023 mmol/L) (24:76, v/v), and the flow rate was 1.0 mL/min [56].

Photolysis products of tigecycline from quantum chemical calculations were analyzed via high-resolution mass spectrometry (HRMS, Xevo G2-XS QToF, Waters, USA). The binary mobile phase was 90% CH_3_OH (0.1% methanoic acid). The ionization and desolvation parameters were optimized as follows: voltage, 3.5 kV; ion source temperature 110 °C, desolvation temperature 400 °C; and nitrogen gas flow rate, 800 L·h^-1^.

### 5.9 Quantum chemical calculation

All calculations in this study were performed using the Gaussian 16 program package [57]. Full geometry optimizations were performed to locate all stationary points, using the B3LYP method with the def2svp basis [58], namely, B3LYP/def2svp. Dispersion corrections were computed using Grimme’s D3(BJ) method in optimization [59]. Harmonic vibration was performed at the same frequency to guarantee that there was no imaginary frequency in the molecules, i.e., they were located on the minima of the potential energy surface. Convergence parameters of the default threshold were retained (maximum force within 4.5×10^−4^ Hartrees/Bohr and root mean square (RMS) force within 3.0×10^-4^ Hartrees/Radian) to obtain the optimized structure. The optimal structure was identified, given that all calculations for structural optimization converged within the convergence threshold of no imaginary frequency during the vibration analysis. The solvation energies were calculated from previous studies [60].

### 5.10 Electron paramagnetic resonance (EPR) spectroscopy

The EPR analysis [61–63] involved two spin-trapping agents, DMPO (5,5-dimethyl-1-pyrroline-N-oxide) and TEMP (2,2,6,6-tetramethylpiperidine), to detect transient free radicals of tigecycline with/without Cu^2+^. For DMPO, hydroxyl radicals (•OH) were generated via a fenton reaction and incubated with 100 mM DMPO at 25°C for 5 min and 30 min to form stable spin adducts. TEMP was reacted with singlet oxygen (^1^O_2_) under visible light irradiation and oxygenated conditions. Triplicate measurements ensured reproducibility, and control experiments (in the absence of spin traps/precursors) confirmed signal specificity.

### 5.11 Biofilm formation and inhibition assays

The tested bacteria were treated with different concentrations of tigecycline or CuSO_4_ in TSB/LB medium for 24 h at 37 °C to form biofilms. The supernatant was subsequently removed, and the biofilms were washed with PBS and stained with 0.1% crystal violet (CV). The CV-stained biomass was solubilized with 30% glacial acetic acid and quantified by measuring the absorption values at OD_570 nm_. Simultaneously, quantitative analysis of bacterial colonization within the biofilm was conducted via CFUs enumeration.

### 5.12 Leakage of nucleic acid and protein

Following differential treatments, the bacterial cultures were centrifuged at 12,000×*g* for 15 min, and the resulting supernatants were aseptically harvested for subsequent tests. Each supernatant was measured at 280 nm or 260 nm to evaluate the bacterial leakage of protein or nucleic acid, respectively.

### 5.13 Intracellular ATP, ROS, Fe^2+^ contents and inner membrane (IM) permeability assay

Bacteria in the mid-logarithmic phase were treated with different concentrations of tigecycline or CuSO_4_ for 6 h at 37 °C. Then, the cultures were collected, centrifuged, and preserved in suspension. The intracellular ATP concentrations were quantified using an enhanced ATP assay kit (Beyotime, Shanghai, China) based on a standardized calibration curve, strictly following the manufacturer’s analytical protocols. ROS levels were assessed using a fluorometric detection system (DCFH-DA, 2’,7’-dichlorodihydrofluorescein diacetate, MEDCHEMEXPRESS LLC, China), adopting equivalent methodological rigor to ensure inter-assay consistency. Additionally, the bacteria were probed with 10 nM propidium iodide (PI) with the indicated concentrations of drugs, and the fluorescence intensity was measured at an excitation wavelength of 535 nm and an emission wavelength of 615 nm. The Fe^2+^ content was assessed using an Fe^2+^ colorimetric assay kit (Elabscience Biotechnology Co., Ltd., China) [64].

Bacterial suspensions were cotreated with 10 nM PI and specified agents, followed by spectrofluorometric quantification of fluorescence intensity using standardized parameters (λ_ex_= 535 nm, λ_em_= 615 nm) with a microplate reader [54].

### 5.14 Transcriptomics and RT-PCR analysis

To elucidate the molecular mechanisms underlying bacterial regulation by different concentrations of Cu^2+^, experimental groups were established: low-concentration Cu^2+^ (32 μg/mL), high-concentration Cu^2+^ (256 μg/mL), tigecycline (1/8 μg/mL), and a control. Transcriptomic and RT-qPCR analyses were performed across *S. aureus* USA300 and *E. coli* ZJ487, focusing on pathways associated with membrane transport systems, to identify critical proteins targeted by non-bactericidal Cu^2+^ for bacterial survival and proliferation. Transcriptome data were collected by Novogene Co., Ltd. (Beijing, China) [65–67].

### 5.15 Glue strip qualitative proteome

Overnight cultured bacteria were treated with different concentrations of CuSO_4_ for 6 h, 12 h, or 24 h at 37 °C. Bacterial cells were harvested, lysed, and subjected to protein separation via SDS-PAGE. Differential protein bands were excised for qualitative proteomic analysis. The identified proteins were confirmed through database interrogation with rigorous quality control of mass spectrometry data. Functional annotation was performed using the Cluster of Orthologous Groups of proteins (COG), Gene Ontology (GO), KEGG pathway, and InterProScan (IPR) annotation databases.

### 5.16 Surface motility of *S. typhimurium*

Overnight cultured *Acinetobacter baumannii* (*A. baumannii*) ATCC17978 was diluted to obtain an OD_600_ of 0.5. Next, 3 µL of prepared bacteria was dropped on 0.3% agar plates containing different concentrations of CuSO_4_ (0, 16, or 64 µg/mL) and FeC_6_H_5_O_7_ (0, 100, or 500 µg/mL). After static culture at 37 °C for 12 h, the *A. baumannii* ATCC17978 on agar plates was photographed, and its mobility diameter was measured [68, 69].

### 5.17 Safety evaluation and cytoprotection assessment

The potential toxicity of Cu^2+^ and tigecycline (light and dark) to the A549 cells was determined using a Cytotoxicity Detection Kit (LDH) (Roche, Basel, Switzerland) after coincubation for 12 h. Cells treated with Triton X-10 (0.2%) were used as a positive control, and untreated cells were used as a negative control.

The invasion ability of *E. coli* ZJ487 in RAW264.7 cells was monitored using a verified gentamicin protection assay. The cells were cultured overnight and treated with tigecycline (1/4 μg/mL), FeC_6_H_5_O_7_ (16 μg/mL), tigecycline + FeC_6_H_5_O_7_ (1/4 μg/mL + 16 μg/mL), or the blank control (0 μg/mL + 0 μg/mL) and were infected with *S. aureus* USA300 and *E. coli* ZJ487 at an MOI (10:1) for 2 h, 6 h, or 12 h. Then, the cells were washed, and bacteria that had not been affected by gentamicin were removed. The bacterial cells were lysed, serially diluted in PBS, and plated on LB agar for CFUs enumeration following 24-h incubation at 37 °C.

### 5.18 *G. mellonella* larvae infection model

A total of 1×10^4^ to 1×10^7^ CFUs of prepared *E. coli* ZJ487 were microinjected through the left pro-leg to establish the optimal bacterial challenge dosage. The toxicity of high-dose Cu(C_6_H_11_O_7_)_2_ to *G. mellonella* larvae was also evaluated. After determining the optimal modeling parameters, an infection model was established. At 2 h post-infection, the larvae were assigned to six experimental cohorts: the Cu(C_6_H_11_O_7_)_2_ monotherapy (2 mg/kg), tigecycline monotherapy (0.2 mg/kg), combination therapy Cu(C_6_H_11_O_7_)_2_: tigecycline at 1:1, 1:5, and 1:10), and only infected groups. The infected larvae were maintained under light conditions and monitored for 7 days to evaluate their therapeutic efficacy against *E. coli* ZJ487 infection. Mortality was defined as larval unresponsiveness to mechanical stimulation accompanied by melanization [70, 71].

To further investigate bacterial clearance dynamics, a sublethal-dose infection model was implemented. Larvae were euthanized at 48 h post-therapeutic intervention for quantitative assessment of bacterial colonization through CFUs enumeration.

### 5.19 Murine skin infection model

The mice were randomly divided into 5 groups (total = 70) and anesthetized with intraperitoneal tribromoethanol. The backs of the mice were then shaved, and 10 mm diameter wounds were made via surgical punches. Then, *S. aureus* USA300 was resuspended at a concentration of 3 × 10^8^ CFUs/mL (50 μL), and were subcutaneously injected into the shaved area [70, 72]. At 2 h after infection, different combinations of Cu(C_6_H_11_O_7_)_2_ and tigecycline were dripped on near the infected skin. The appearance of the skin under the naked eye and microscope was monitored for 15 days (n = 6 per group), body weight was measured daily for 20 days (n = 3 in each group were selected for weight measurement), and the wounded skin was aseptically excised, and homogenized to count bacterial CFUs in the skin (n = 8 per group).

### 5.20 Statistical analysis

Statistical analyses were conducted using GraphPad Prism, and the data are presented as the mean ± standard deviation (SD). Sample sizes (n) are explicitly stated in the respective figure legends. The statistical significance was analyzed by the unpaired Student’s t-test method and one-way ANOVA to calculate *P*-values. The least significant difference test was performed, and the differences were considered to be statistically significant at *p* < 0.05.

## Supporting Information

The Supporting Information is available free of charge via the Interne. (Supplementary materials).

## Acknowledgments

We are grateful to Professor Wang Yang from China Agricultural University for providing us with the clinical isolates. We also thank all members of the Deng Lab for the helpful discussions. This work was supported by the National Natural Science Foundation of China (grant 32560874, 32202856 and U23A20242).

## Ethics Statement

All animal procedures were conducted following protocols approved by the Ethics Review Committee of Animal Experimentation at Ningxia University (No. NXU-2025-062) and in compliance with the principles of the Animal Welfare Act.

## Author contributions

YLZ and JFW designed this project; JJX, QYZ, HRL and KG performed experiments; QHL, YYL and LM analyzed the data and prepared figures; YLZ and JFW provided resource support; YLZ and JJX drafted and revised this manuscript. All authors reviewed, revised, and approved the final report.

## Data and materials availability

All data to evaluate the conclusions in the paper are present in the Manuscript and/or the Supplementary Materials. The detailed information for all datasets can be seen in Experimental Section.

---

**Figure S1.**
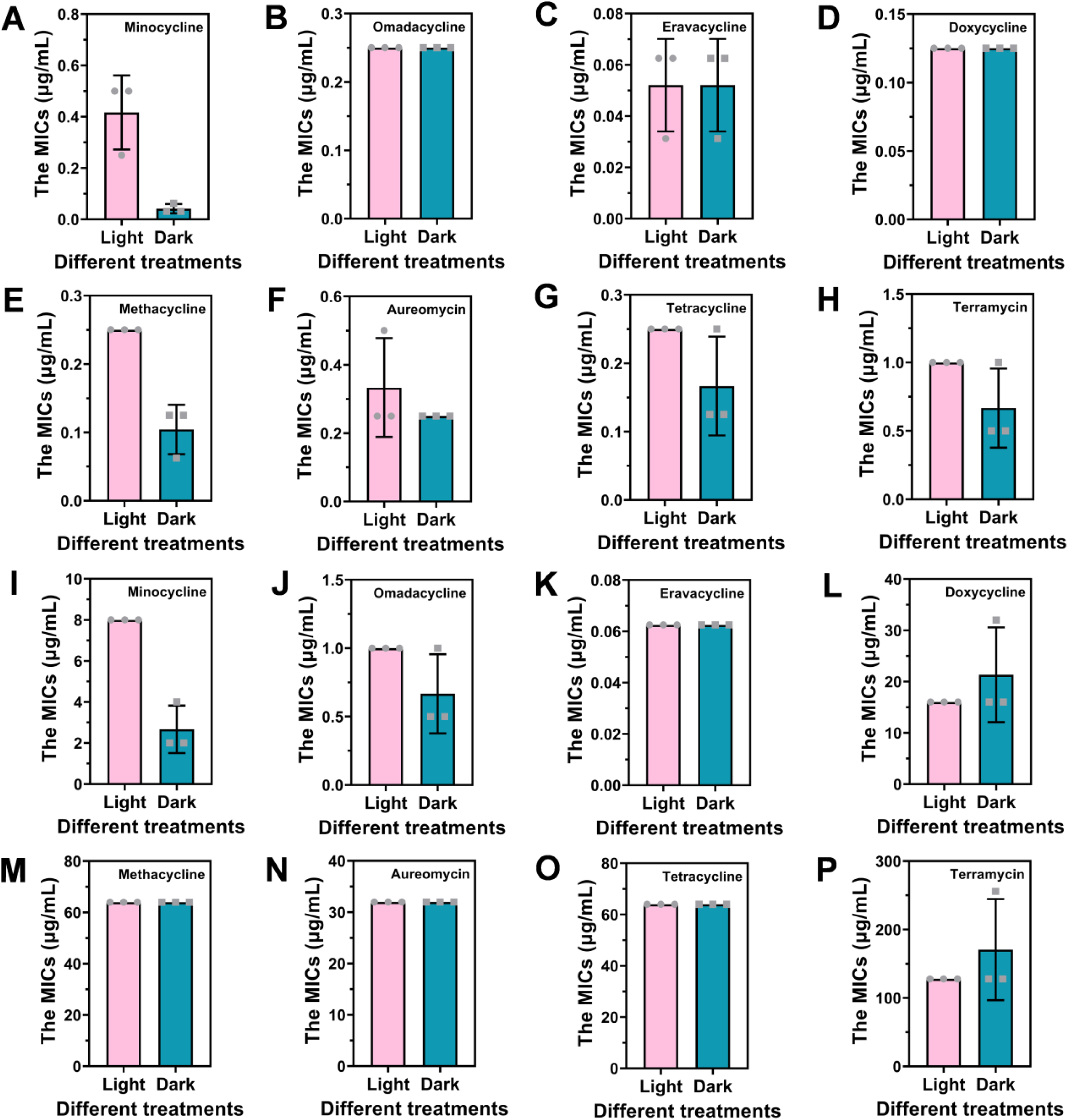
Detection of MIC values of eight tetracyclines against *S. aureus* USA300 and *E. coli* ZJ487 under light and dark conditions. Minocycline (**A**), omacycline (**B**), eravacycline (**C**), doxycycline (**D**), methacycline (**E**), aureomycin (**F**), tetracycline (**G**), and terramycin (**H**) for *S. aureus* USA300. Minocycline (**I**), omacycline (**J**), eravacycline (**K**), doxycycline (**L**), methacycline (**M**), aureomycin (**N**), tetracycline (**O**), and terramycin (**P**) for *E. coli* ZJ487. Three independent experiments were performed (n = 3)

**Figure S2.**
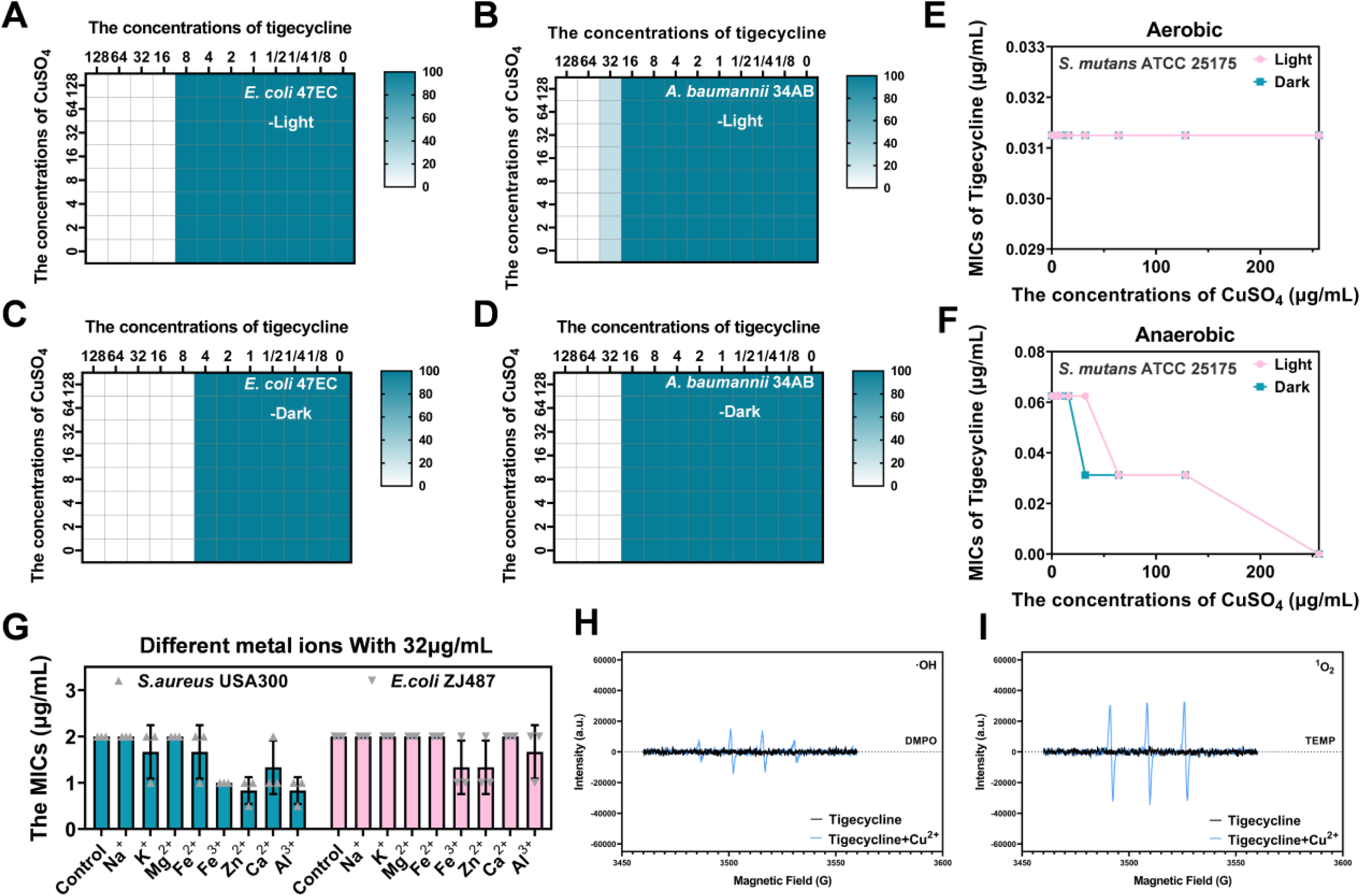
(**A-D**) The MIC values of CuSO_4_ combined with tigecycline against Tet(X3)- and Tet(X4)-positive bacteria (*E. coli* 47EC carrying Tet(X4), *A. baumannii* 34AB carrying Tet(X3)) under light and dark conditions were detected by using the checkerboard method. (**E, F**) The MIC values of CuSO_4_ combined with tigecycline against *S. mutans* ATCC25175 under light and dark conditions, and under both aerobic and anaerobic conditions. (**G**) The MIC values of tigecycline with different metal ions (32 μg/mL) against *S. aureus* USA300 and *E. coli* ZJ487 under light conditions. EPR characterization of hydroxyl radicals (·OH) (**H**) and singlet oxygen (^1^O_2_) (**I**) across experimental groups after treatment for 5 min under light conditions.

**Figure S3.**
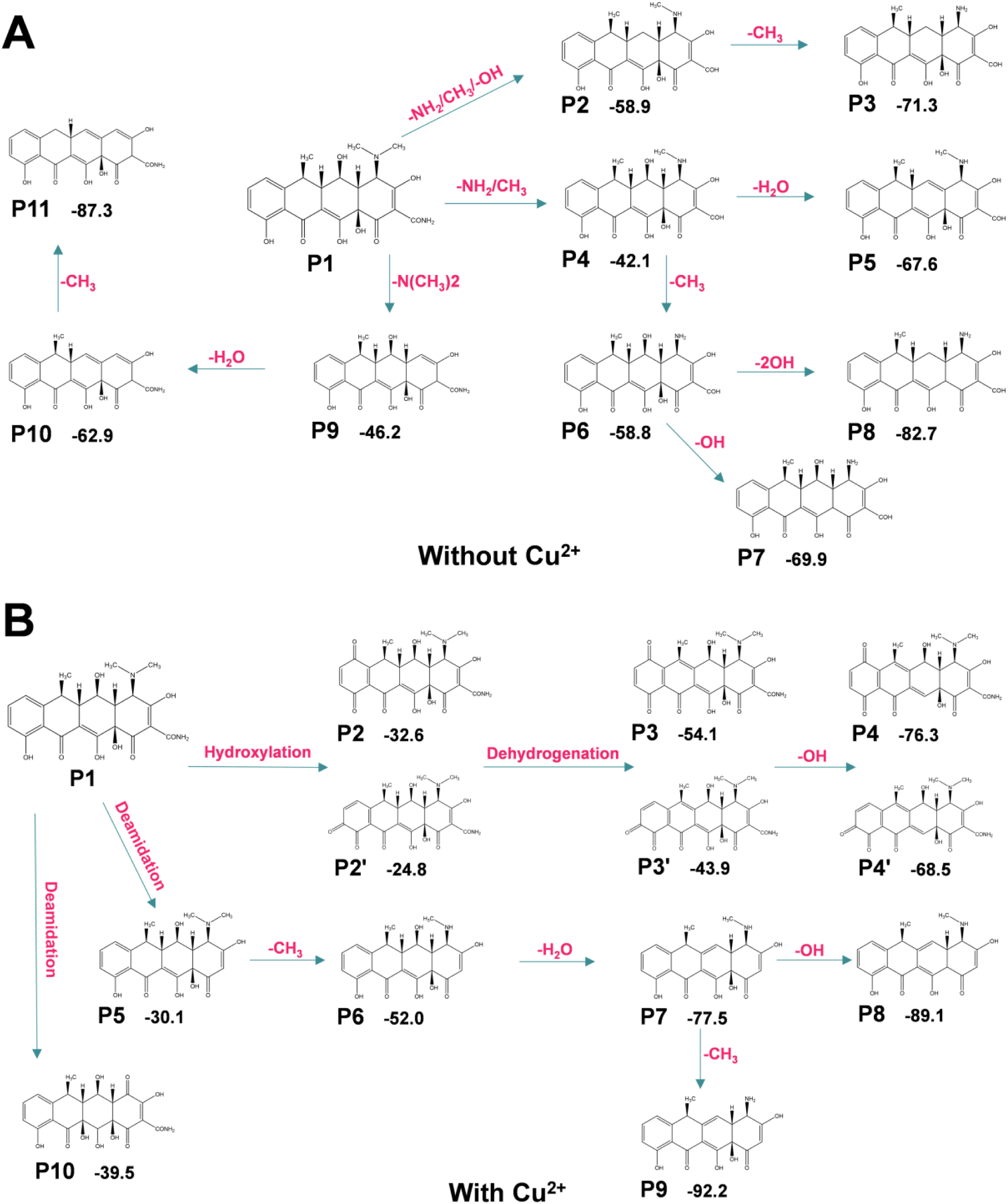
Photolysis products and degradation pathways of doxycycline with (B) or without (A) CuSO_4_ under light irradiation.

**Figure S4.**
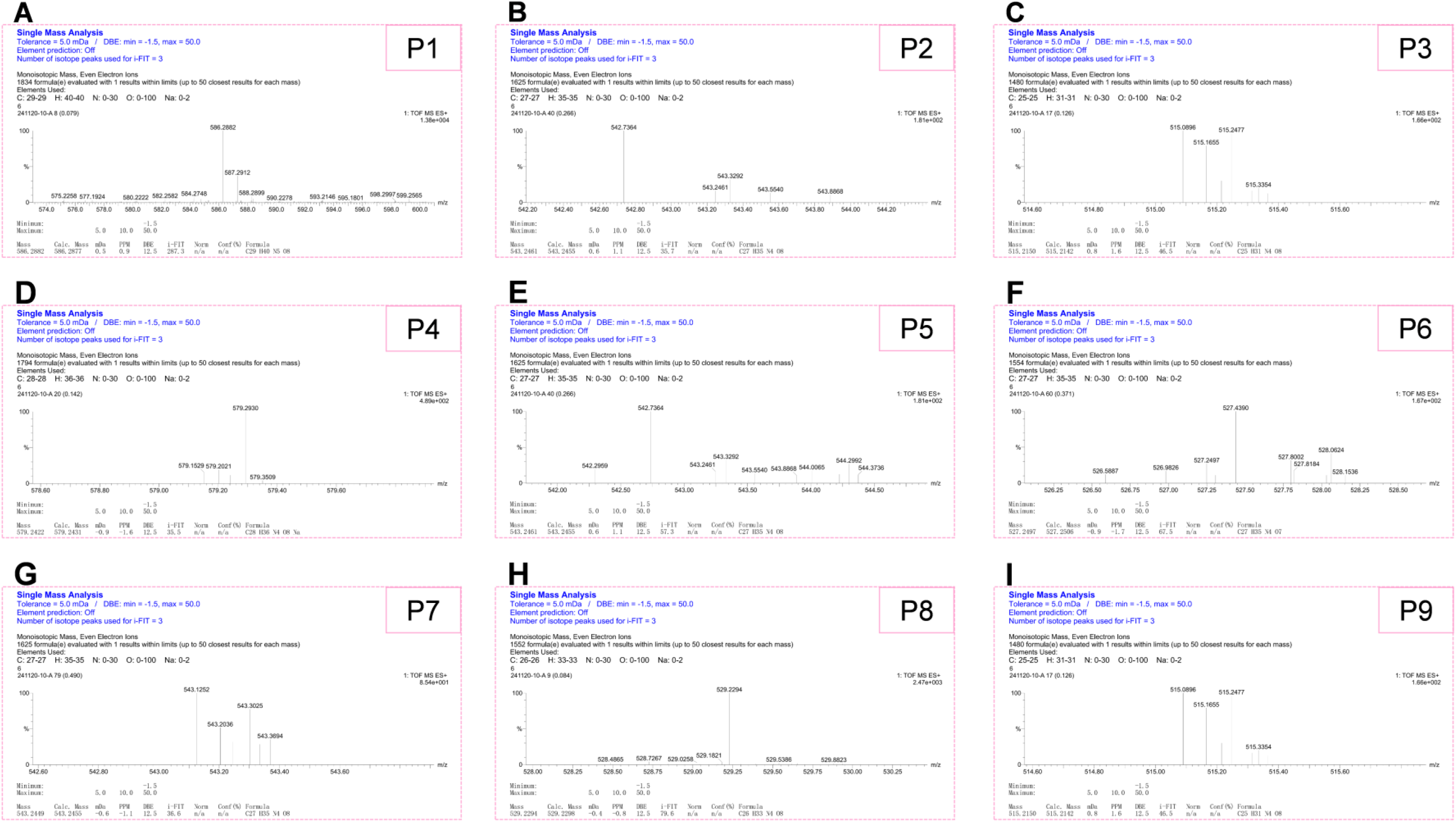
Single mass analysis of the photolysis product of tigecycline without Cu^2+^. (**A** – **I**) correspond to nine photolysis products.

**Figure S5.**
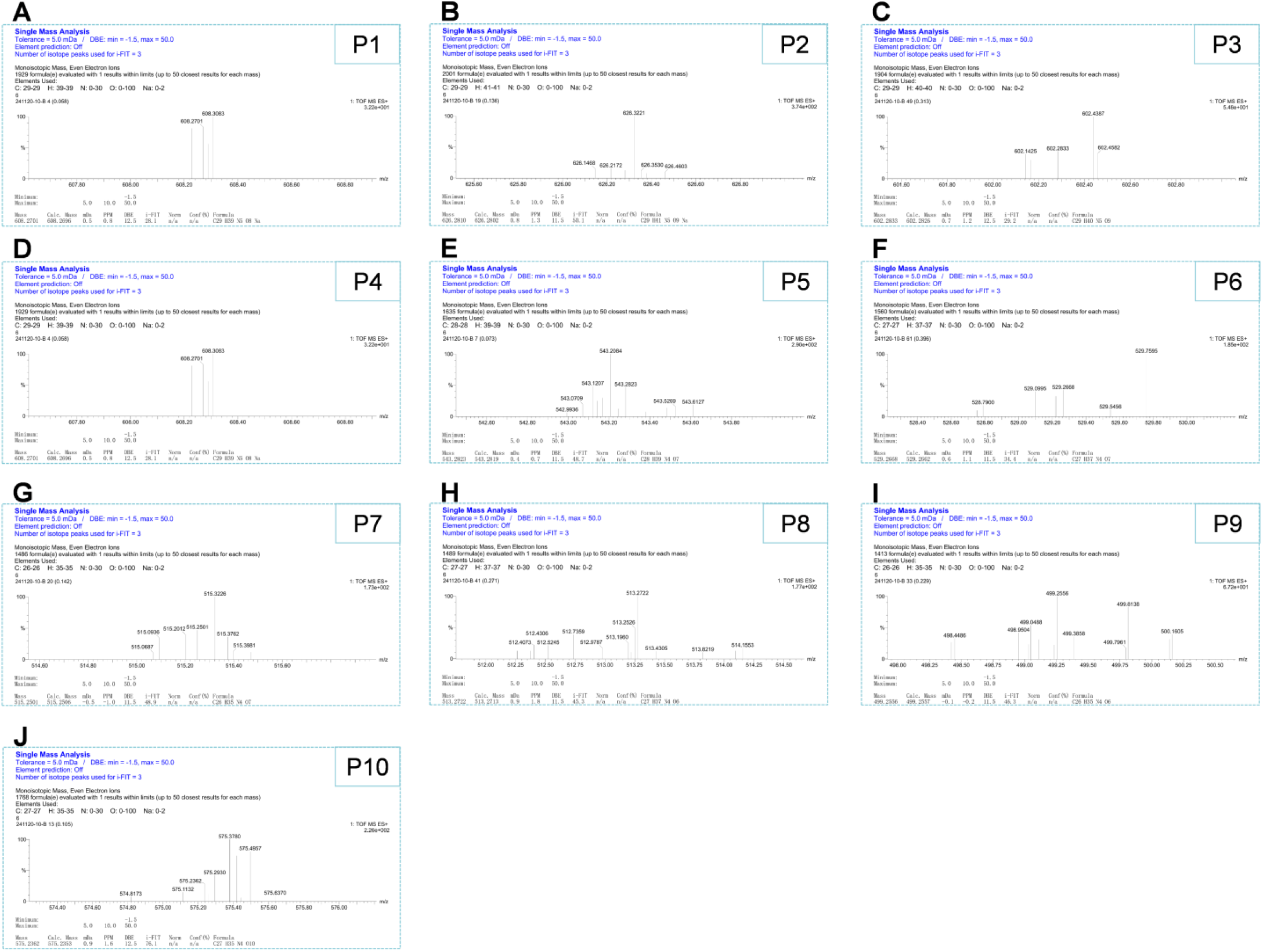
Single mass analysis of the products of the photolysis of tigecycline with Cu^2+^. (**A** – **J**) correspond to ten photolysis products.

**Figure S6.**
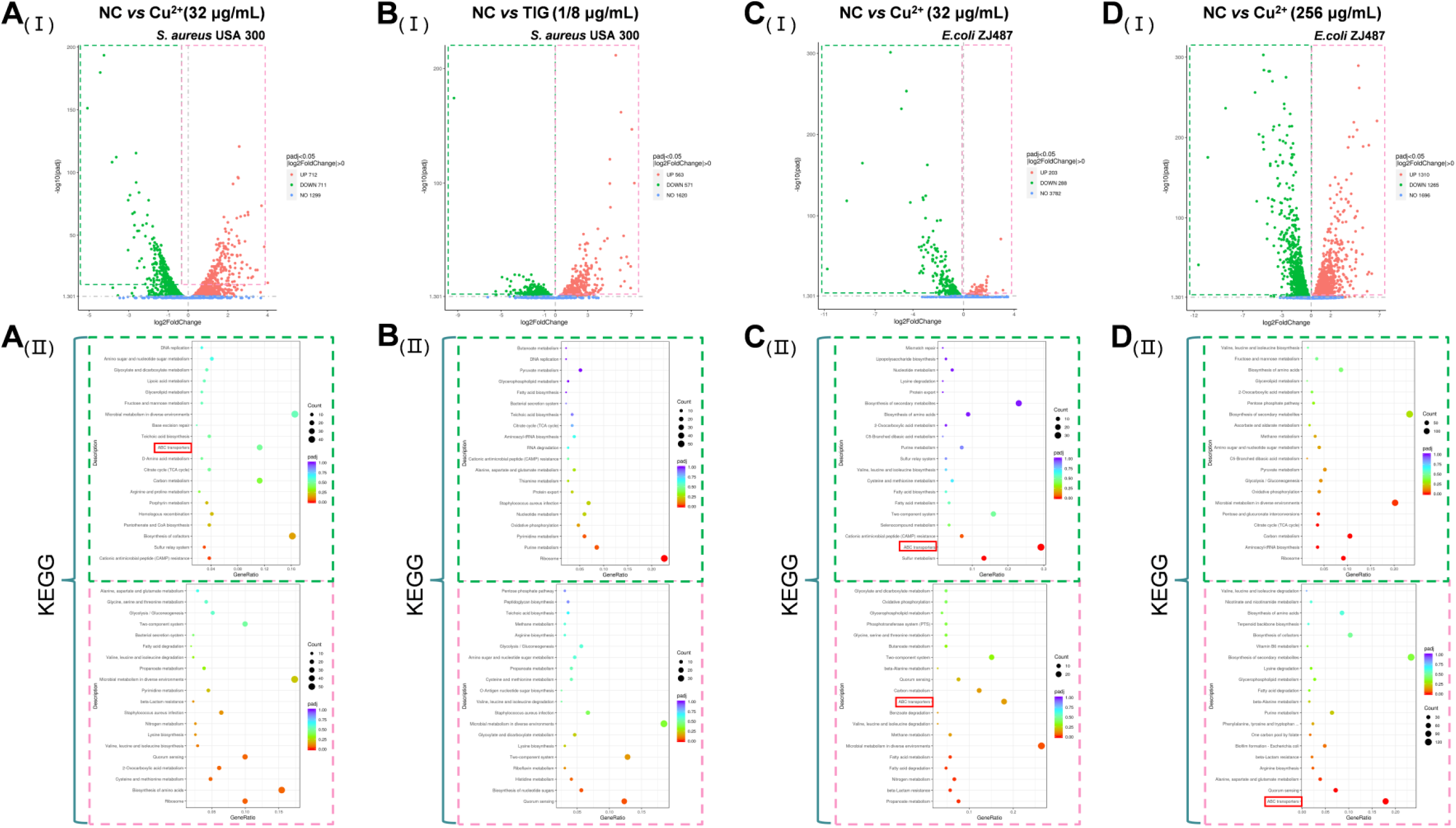
Transcriptomic analysis. **A_(Ⅰ)_**-**D_(Ⅰ)_**-The differentially expressed genes of *S. aureus* USA300 and *E. coli* ZJ487 treated with CuSO_4_, tigecycline or control. **A_(Ⅱ)_**-**D_(Ⅱ)_** Analysis of KEGG pathway enrichment analysis of genes that were differentially expressed between different groups, and the mutually downregulated genes were focused on the ABC transporters of bacteria (outlined in red).

**Figure S7.**
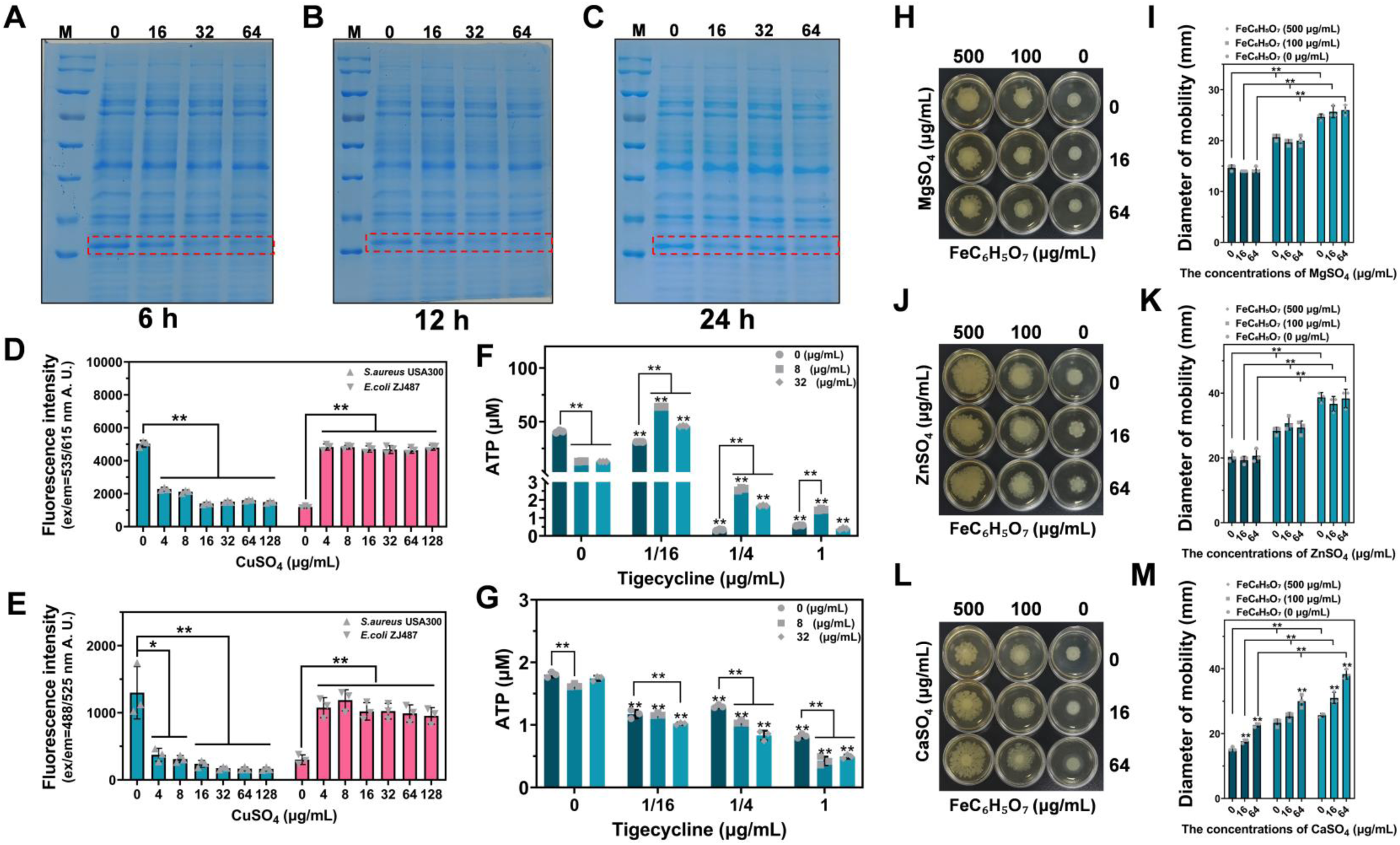
Assessment of the effects of Cu^2+^-mediated modulation on bacterial cellular integrity, biofilm formation, and surface motility. All protein samples collected from *E. coli* ZJ487 treated with different concentrations of CuSO_4_ after 6 h (**A**), 12 h (**B**), and 24 h (**C**), were loaded on SDS-PAGE gels and analyzed via Coomassie brilliant blue staining. Enhanced ROS accumulation and the fluorescence value of PI in *E. coli* ZJ487 cultured with CuSO_4_. (**D, E**) Exposure to CuSO_4_ increased ROS generation and PI influx in *E. coli* ZJ487, whereas *S. aureus* USA300 exhibited paradoxical attenuation. Detection of intracellular ATP levels in *S. aureus* USA300 (**F**) and *E. coli* ZJ487 (**G**). MgSO_4_ and ZnSO_4_ had no inhibitory effects on surface motility in *A. baumannii* ATCC 17978 (**H-K**), and CaSO_4_ even promoted the surface motility of *A. baumannii* ATCC 17978 (**L**, **M**). Experiments dates were shown as mean±SD. * indicates *P* < 0.05. ** indicates *P* < 0.01

**Figure S8.**
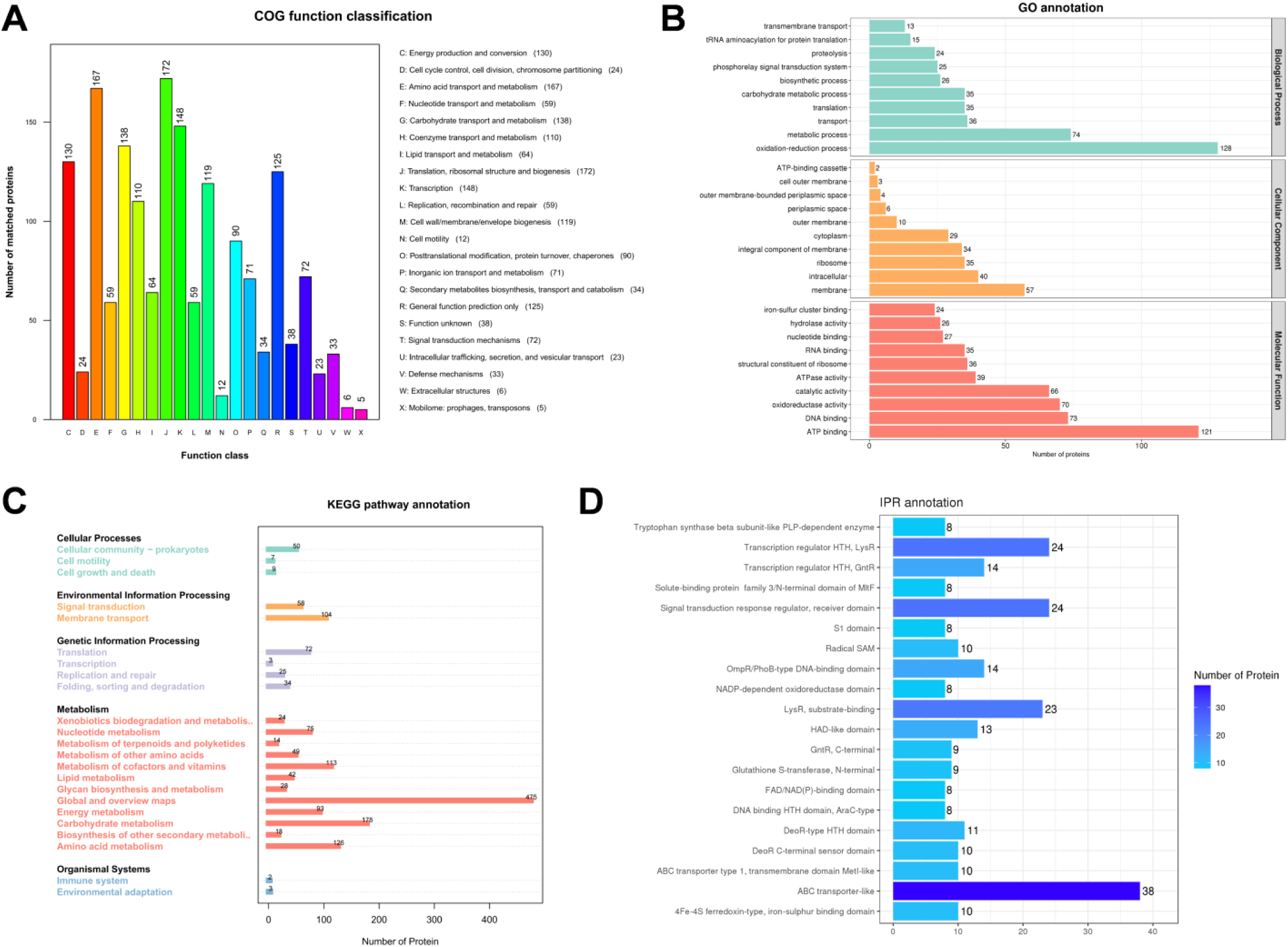
Interaction spectrum qualitative proteome analysis. Integrated functional annotation analyses using COG function classification (**A**), GO annotation (**B**), KEGG pathway annotation (**C**), and IPR annotation (**D**) databases culminated in the identification of a high-potential target.

**Figure S9.**
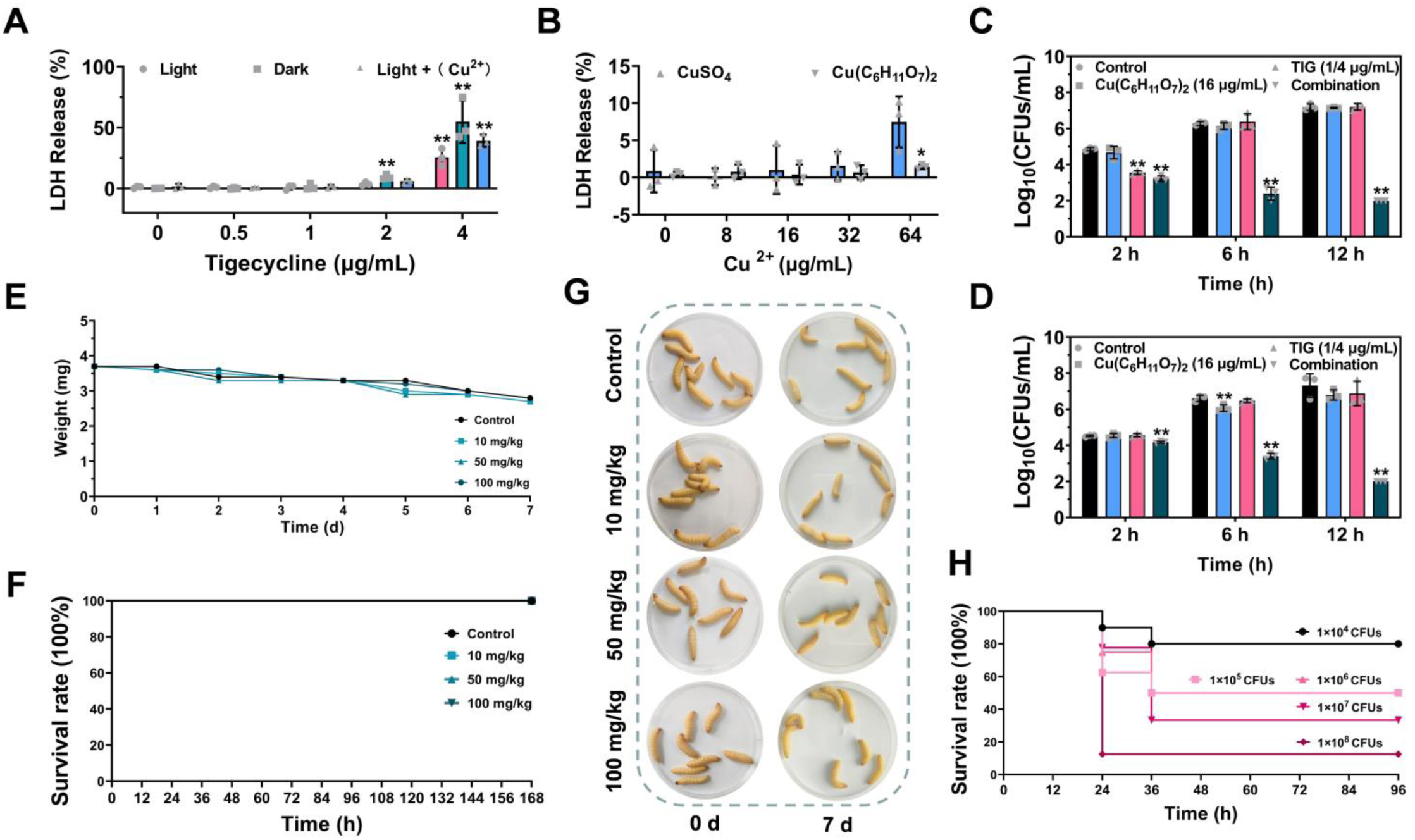
Preparatory tests were performed before conducting formal trials. (**A, B**) LDH release after different treatments. The invasion of *S. aureus* USA300 (**C**) and *E. coli* ZJ487 (**D**) in RAW264.7 cells. After being treated with copper gluconate at concentrations of 0 mg/kg, 10 mg/kg, 50 mg/kg and 100 mg/kg, the body weight (**E**) and survival (**F, G**) of the *G. mellonella* larvae were monitored during the 7-day observation period (n = 8). (**H**) Survival of *G. mellonella* larvae after infection with different concentrations of *E. coli* ZJ487was recorded (n = 10). Experiments dates were shown as mean±SD; * indicates *P* < 0.05. ** indicates *P* < 0.01.

**Table S1.** The information of the tested bacteria in this study.

| Species | Source | Drug resistance description |
| --- | --- | --- |
| *S.aureus* USA300 | Laboratory strain | Methicillin-Resistant *staphylococcus aureus* |
| *E. coli* ZJ487 | Human intra-abdominal fluid<sup>1</sup> | Carried *mcr-1* gene and *ndm-1* gene |
| *S.aureus* USA400 | Laboratory strain | Methicillin-Resistant *staphylococcus aureus* |
| *S.aureus* ATCC25904 | Laboratory strain | \ |
| *S.aureus* ATCC29213 | Laboratory strain | Sensitive to oxacillin |
| *S.aureus* 8325-4 | Laboratory strain | Methicillin-Resistant *staphylococcus aureus* |
| *S. aureus* ST2032 | Animal farm<sup>2</sup> | Methicillin-Resistant *staphylococcus aureus* |
| *S.aureus* T144 | Animal farm<sup>3</sup> | Methicillin-Resistant *staphylococcus aureus*<br>Hypervirulent model strain, isolated from pigs |
| *S.aureus* JL02 | A-Class Hospital<sup>2</sup> | Methicillin-Resistant *staphylococcus aureus* |
| *L. monocytogenes* EGD | Laboratory strain | \ |
| *S. pneumoniae* D39 | Laboratory strain | \ |
| *E. faecalis* NS1 | Animal farm | Macrolide resistance |
| *E. coli* J53 | Laboratory strain | Recipient of drug resistance gene |
| *E. coli* ATCC25922 | Laboratory strain | \ |
| *E. coli* DH5α+pAM401-TEM-1 | Recipient of *tem-1* gene | Carried a *tem-1* gene |
| *E. coli* D3 | Animal farm<sup>4</sup> | Carried a *ndm-1* gene |
| *E. coli* ZC-YN7 | Animal farm<sup>4</sup> | Carried a *ndm-9* gene |
| *A. baumannii* ATCC19606 | Laboratory strain | \ |
| *A. baumannii* ATCC17978 | Laboratory strain | \ |
| *S. typhimurium* ATCC14028 | Laboratory strain | \ |
| *S. typhimurium* HYM2 | Animal farm<sup>5</sup> | Carried a *mcr-1* gene |
| *S. typhimurium* SL1344 | Derived from the virulent strain SL1344<sup>6</sup> | \ |
| *K. pneumoniae* ATCC700603 | Laboratory strain | \ |
| *K. pneumoniae* ZJ02 | Remote tertiary care hospital<sup>1</sup> | Carried a *mcr-1* gene |
| *P. aeruginosa* ATCC27853 | Laboratory strain | \ |
| *E. coli* 47EC | Animal farm<sup>7</sup> | Donor of tet(X3)-carrying plasmid p34AB (pig) |
| *A. baumannii* 34AB | Animal farm<sup>7</sup> | Donor of tet(X4)-carrying plasmid p47EC (pig) |
| *S. mutans* ATCC25175 | Laboratory strain | \ |

